# Single-cell chromatin landscape and DNA methylation patterns reveal shared molecular programs in human tumor and non-tumor tissue CCR8^+^ Treg cells

**DOI:** 10.1101/2025.05.14.653926

**Authors:** Kathrin Luise Braband, Tamara Kaufmann, Niklas Beumer, Malte Simon, Sara Salome Helbich, Delia Mihaela Mihoc, Morten Michael Voss, Annekathrin Silvia Nedwed, Katharina Bauer, Jan-Philipp Mallm, Tanja Ziesmann, Ute Distler, Stefan Tenzer, Christoph Eckert, Michael Volkmar, Dieter Weichenhan, Marion Bähr, Christoph Plass, Matthias Linke, Stefan Diederich, Matthias Klein, Tobias Bopp, Hansjoerg Schild, Andreas Henkel, Edoardo Filippi, Federico Marini, Benedikt Brors, Matthias Martin Gaida, Michael Delacher

**Affiliations:** Institute of Immunology, University Medical Center Mainz, 55131 Mainz, Germany; Research Center for Immunotherapy, University Medical Center Mainz, 55131 Mainz, Germany; Division of Applied Bioinformatics, German Cancer Research Center (DKFZ), 69120 Heidelberg, Germany; Evotec SE, 22419 Hamburg, Germany; Institute of Medical Biostatistics, Epidemiology and Informatics (IMBEI), University Medical Center Mainz, 55131 Mainz, Germany; Single Cell Open Lab, German Cancer Research Center (DKFZ) and Bioquant, Heidelberg, Germany; German Cancer Research Center (DKFZ), INF 280, 69120, Heidelberg, Germany; Helmholtz Institute for Translational Oncology Mainz (HI-TRON Mainz) - A Helmholtz Institute of the DKFZ, Obere Zahlbacherstr. 63, 55131 Mainz, Germany; Institute of Pathology, University Medical Center Mainz, 55131 Mainz, Germany; Division of Cancer Epigenomics, German Cancer Research Center (DKFZ), INF 280, 69120, Heidelberg, Germany; Institute of Human Genetics, University Medical Center Mainz, 55131 Mainz, Germany; German Cancer Consortium (DKTK), German Cancer Research Center (DKFZ), Im Neuenheimer Feld 280, 69120 Heidelberg, Germany; High Performance Computing and Data Center, University of Mainz, 55128 Mainz, Germany; Department of Nephrology and Rheumatology, Center of Immunotherapy, University Medical Center Mainz, 55131 Mainz, Germany; National Center for Tumor Diseases (NCT), Im Neuenheimer Feld 460, 69120 Heidelberg, Germany; Faculty of Biosciences, Heidelberg University, 69120 Heidelberg, Germany; Faculty of Medicine, Heidelberg University, 69120 Heidelberg, Germany; TRON, Translational Oncology at the University Medical Center, JGU-Mainz, 55131 Mainz, Germany

## Abstract

Regulatory T (Treg) cells, a subset of CD4^+^ T cells, play a crucial role in immunoregulation. Notably, CCR8-expressing Treg cells in tissues also contribute to organ homeostasis and repair. To determine whether these tissue-regenerative programs are active in the tumor microenvironment, we employed single-cell chromatin accessibility and genome-wide DNA methylation analyses to investigate CCR8^+^ tissue Treg cells isolated from human tumor and adjacent tumor-free tissues. Our findings indicate that CCR8^+^ tissue Treg cells from tumor and corresponding tumor-free tissues exhibit a high degree of similarity, suggesting that the tumor microenvironment may harbor highly activated tissue Treg cells. This observation was consistent across various tumor types and origins, including primary tumors and metastases. Using quantitative proteomics, we identified several candidate factors associated with the regenerative and suppressive programs of Treg cells, which may serve as potential reservoir of druggable targets for future therapeutic interventions.

## INTRODUCTION

Regulatory T (Treg) cells, a subset of CD4^+^ T cells, play a crucial role in immunoregulation, including self-tolerance and the down-modulation of inflammation after pathogen clearance^1^. In addition to their immunoregulatory functions, Treg cells have been shown to promote organ homeostasis and regeneration in mice. This tissue-regenerative program is characterized by shared phenotypes between tissues, transient multi-tissue migration patterns, and common molecular dependencies under steady-state conditions^2^.

Tissue Treg cells have been shown to promote tissue regeneration in a variety of tissues in murine model systems; these include the skin, where they promote hair follicle stem cell differentiation in the steady state^3^ and promote tissue repair upon UVB-induced skin damage^4^ or mechanical injury^5^; the liver, where they contribute to homeostasis by regulating bile acid synthesis^6^; the visceral adipose tissue, where they exert functions in metabolic control^7^; the kidney, where they promote tissue repair after injury^8^; the lungs, where they also play a role in tissue regeneration and the reduction of fibrosis^9, 10, 11, 12, 13, 14^; the skeletal muscle and the heart, where they have regenerative functions in dystrophic muscle^15, 16^ and after myocardial infarction^17, 18, 19^, respectively; and the CNS, where they have been shown to limit brain damage after ischemic stroke^20, 21^ and promote oligodendrocyte differentiation and re-myelination in a mouse model of multiple sclerosis^22^.

All these findings indicate an important role of tissue Treg cells for tissue homeostasis and repair, but they also raise the question of whether Treg cells exhibit tissue repair or -remodeling functions in the tumor microenvironment (TME). Recent studies have suggested that Treg cells in the TME may promote tumor growth and progression^23, 24, 25, 26^. In the murine system, it is known that the EGFR-ligand amphiregulin (Areg) plays a central role in tissue repair^27^ and has detrimental effects in a variety of tumor models: in a lung tumor model, Areg deficiency in Treg cells led to delayed tumor progression^23^, and Areg produced by Treg cells has further been shown to actively promote tumor cell proliferation, epithelial-mesenchymal transition, and the formation of metastases^24, 25^. Areg has been found to induce PD-L1 expression in cancer cells, promoting an immunosuppressive TME^26^. Together, these studies suggest that the tissue regeneration program of murine tissue Treg cells adds to their pro-tumorigenic function in the TME.

In humans, however, shared molecular programs between Treg cells isolated from tumors and normal tissues have not yet been investigated in detail, and human Treg-derived factors that might promote tissue regeneration are yet to be determined. The ontogeny and precise classification of human tumor-resident Treg cells is still under debate, and many efforts have been made to find druggable targets in tumor-resident Treg cells. However, Treg cells isolated from the tumor are often compared with Treg cells isolated from peripheral blood^28, 29, 30, 31, 32, 33, 34^, which, in turn, do not reflect the tissue program^35^. As a result, differences arising from the comparison of tumor versus non-tumor tissue cannot be decoupled from differences arising from the comparison of lymphoid versus non-lymphoid tissue Treg cells. Since Treg cells in solid tumor tissues are the target of many recent clinical trials (NCT05537740, NCT05635643, NCT05518045, NCT06387628, NCT05007782, NCT05101070, NCT05935098, NCT06131398), it is of paramount importance to compare tumor Treg cells to Treg cells isolated from the cognate normal tissue in detail to avoid off-target effects, for example the elimination of the tissue Treg pool in unaffected tissues and organs that could lead to disturbed organ homeostasis and even autoimmune events^33^.

Here, we analyze T cells isolated from different human tumors and the corresponding normal tissue adjacent to the tumor (NAT) using single cell chromatin accessibility sequencing (scATAC-seq), and compare molecular dependencies amongst tissue Treg cells isolated from healthy and malignant tissue for several tumor entities. In addition, we generated tagmentation-based whole-genome bisulfite sequencing (WGBS) datasets^36, 37^ of tissue Treg cells from tumor and NAT, and compared the epigenetic programs with data derived from healthy donor fat and skin tissue Treg cells in a multi-omic epigenetic analytical approach^38^. We used this epigenetic approach since Treg-specific DNA methylation patterns are considered a stable epigenetic mark that can be inherited through multiple cell divisions and can thus be useful for lineage classification of cells^27, 39, 40^. Furthermore, we investigated the transposable element (TE) landscape, another distinguishing feature of tissue Treg cells^38^, in tumor vs NAT Treg cells.

Our data indicate that, on a global scale, tumor-resident Treg cells are epigenetically closely related to Treg cells from tumor-free adjacent tissues, raising the possibility that tumor Treg cells share their molecular programming with tissue Treg cells and do not constitute their own cellular identity. Inducing this tissue Treg programming in naive Treg cells *in vitro* not only mediated tissue regenerative features, but also promoted tumor spheroid growth, indicating that the tissue regeneration program could also be linked to tumor growth and remodeling in humans. Thus, the hypothesis that tumor-resident tissue Treg cells not only promote immune suppression, but also fuel tumor growth via their regenerative program, demarcates them as important cellular targets in the TME. Using transcriptomics and quantitative secretomics, we identified several candidate factors linked to the regenerative and suppressive program of tissue-like Treg cells, which could constitute a reservoir of druggable targets for therapeutic interventions in the future without systemic elimination of the tissue Treg cell pool.

## RESULTS

### CD45RA^-^CCR8^+^ tissue Treg cells are present in human tumor & cognate NAT

The vast majority of experimental datasets on tissue Treg cells come from the murine system. However, a recent multi-omics study has provided insights into human tissue Treg cell biology, revealing programs governing human tissue Treg cell differentiation and delineating the presence of these programs across molecular levels^38^. Human Treg cells can be identified using Treg-specific proteins such as CD25 or FOXP3, while tissue Treg cells express the intracellular transcription factor BATF and the surface receptor CCR8^35, 38^. Nevertheless, the presence of CCR8^+^ tissue Treg cells in human tissues other than skin and fat, as well as their relative frequency and molecular signatures in the tumor microenvironment, remains unclear.

Therefore, we first set out to investigate whether CCR8^+^ Treg cells can be found in various human tissues derived from either human diseased tissues or their normal adjacent tissue. To this end, both pathologically confirmed tumor and NAT were digested using customized and tissue-optimized digestion protocols, followed by bead-based enrichment of CD45^+^ immune cells and analysis via flow cytometry (**Figure 1a**, left and **Extended Data Figure 1a-d**). In total, we analyzed tumor tissue and NAT from primary liver tumors (n=28), metastasis of distal tumors to the liver (n=13), primary kidney tumors (n=18), primary colon tumors (n=12) and primary lung tumors (n=8, **Figure 1a**, right). In the comparison of tumor tissue vs NAT, we detected significantly elevated CD127^-^CD25^+^ Treg cell frequencies of CD4^+^ T cells in primary liver, liver metastasis, colon and lung tumors, while kidney tumors showed no significant increase (**Figure 1a**, upper right). Next, we classified tissue Treg cells as the CD45RA^-^CCR8^+^ subpopulation of CD127^-^CD25^+^ Treg cells and identified a significantly increased frequency in tumor versus cognate NAT for all tumor types (**Figure 1a**, lower right). In addition, we investigated tumor tissue and NAT from pancreatic ductal adenocarcinomas (n=18), bladder tumors (n=6), rectal tumors (n=4), testicular tumors (n=4), and mamma carcinomas (n=3), and identified both CD127^-^CD25^+^ Treg cells as well as CD45RA^-^CCR8^+^ tissue Treg cells in all tissues (**Extended Data Figure 1e-k**). Only in pancreas, basically no immune cells could be detected in tumor-free tissue (**Extended Data Figure 1f, j-k**), raising the possibility that the tumor breaks the immune privilege of this tissue and recruits, amongst other cell types, also CCR8^+^ tissue Treg cells into the TME^41^.

**Figure 1:**
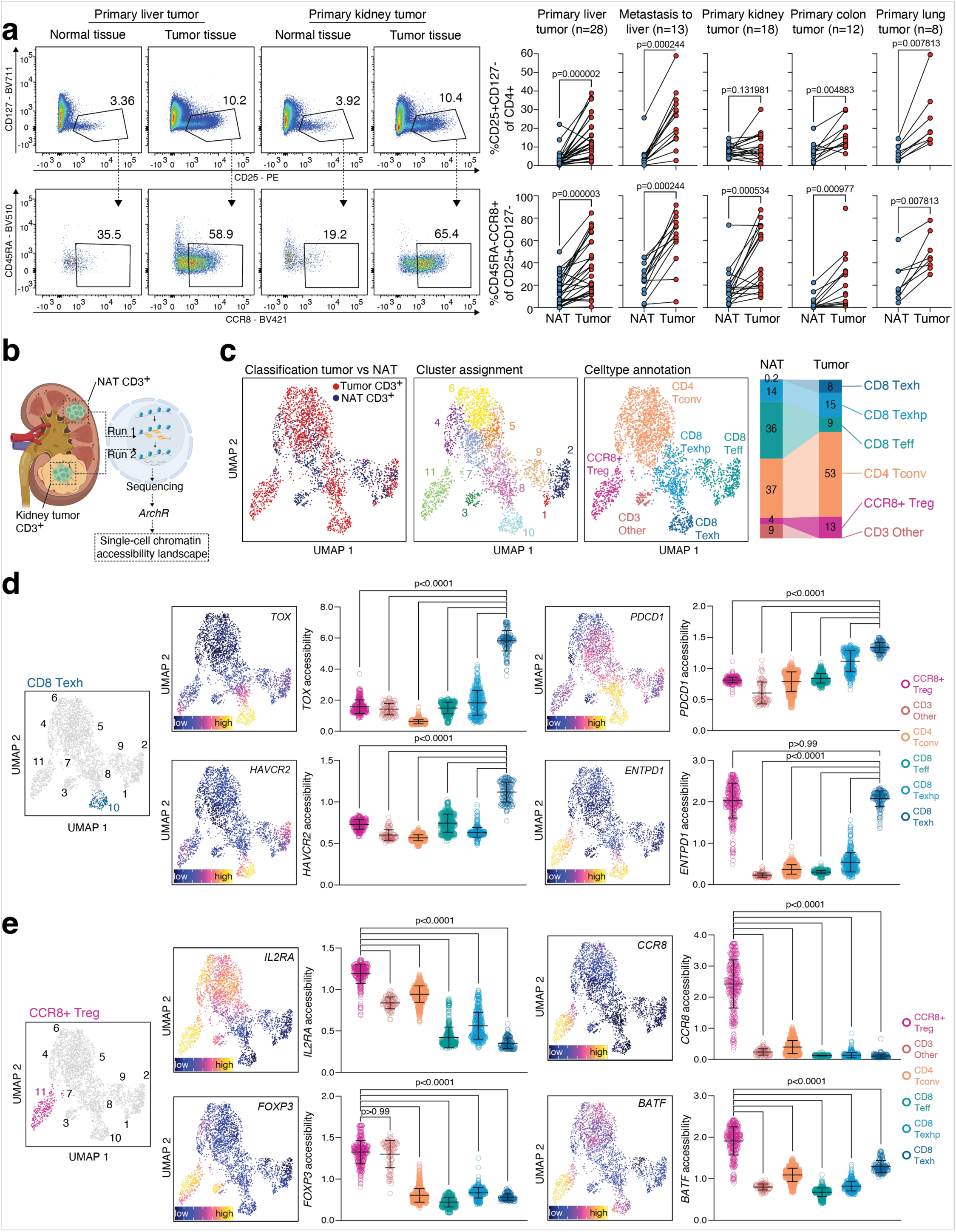
CD45RA^-^CCR8^+^ tissue Treg cells in human tumor and cognate NAT. **(a)** Representative flow cytometry dot plot illustrating presence of Treg cells (CD127^-^CD25^high^ CD4^+^ T cells) and CCR8^+^ Treg cells (CD45RA^-^CCR8^+^ from CD127^-^CD25^high^ Treg cells) in a primary liver and kidney tumor patient. Besides tumor tissue, normal tissue adjacent to tumor (NAT) was analyzed to assess the T cell landscape in tumor-free tissue. Right, Treg cell frequency of CD4^+^ T cells (right, top) and CCR8^+^ Treg cell frequency of Treg cells (right, bottom) for tumor and healthy tissue isolated from human liver (primary tumor n=28, metastases n=13), human kidney (n=18), colon (n=12), and lung (n=8), statistical analysis via Wilcoxon two-tailed matched-pairs signed-rank test. Additional tissues, full gating and representative dot plots in **Extended Data Figure 1**. **(b)** Schematic showing processing of tumor tissue and NAT for scATAC-seq in a single experiment. **(c)** UMAP plot of one exemplary kidney tumor patient (clear cell renal carcinoma) encoding sample information (left), cluster membership (middle), and cell type annotation (right) and stacked bar graph displaying cell type composition (CD3^+^ T cells) in tumor and NAT. **(d-e)** Annotation of CD8 Texh cells by elevated gene activity scores for *TOX*, *HAVCR2*, *PDCD1* and *ENTPD1* and annotation of CCR8^+^ Treg cells by elevated gene activity scores for *IL2RA*, *FOXP3*, *BATF* and *CCR8*. Individual dots illustrate cells, statistical analysis using Kruskal-Wallis and Dunn’s multiple comparisons test. More data related to cell sorting, sample parameters and cluster annotation in **Extended Data Figure 2.** Data representative of several independent experiments with the indicated number of samples (a) or a single experiment with a kidney tumor patient (b-e).

In conclusion, we identified CD45RA^-^CCR8^+^ tissue Treg cells in various human tissues, including kidney, liver, colon, rectum, lungs, bladder, pancreas, and others. Notably, we observed a higher frequency of CD45RA^-^CCR8^+^ tissue Treg cells in the tumor compared to the corresponding normal adjacent tissue. These findings prompt the question of whether tissue Treg cells in tumor and NAT are related, or whether tumor-resident Treg cells represent a distinct and unique cell type.

### Single-cell ATAC-seq identifies CD3^+^ T cell subpopulations in tumor and NAT

To characterize the molecular programs of CD45RA^-^CCR8^+^ tissue Treg cells in an unbiased manner, we employed single-cell chromatin accessibility profiling (scATAC-seq) to analyze both tumor and NAT samples. We digested the tissues using tissue-optimized protocols, followed by fluorescence-activated cell sorting of viable CD3^+^ T cells, nuclei preparation, transposition, and single cell capturing (**Figure 1b**). Both tumor and NAT CD3^+^ T cells were processed using the same reagents and loaded on the same chip to avoid reagent batch effects or handling bias. Sample pre-processing, including quality filtering, as well as downstream analysis and visualization, was performed using *CellRanger ATAC* and *ArchR* ^42, 43^ (**Figure 1c** and **Extended Data Figure 2a-d**). Cell type annotation was performed using gene activity scores of pre-defined marker genes (**Extended Data Figure 2e**). For example, we identified chronically activated and dysfunctional CD8^+^ T cells (“exhausted CD8^+^ T cells”, CD8 Texh) by elevated gene scores for *TOX*, *CTLA4*, *HAVCR2*, *PDCD1*, and *ENTPD1* (**Figure 1d**). A comparison between the TME and NAT showed an increase in CD8 Texh of all CD3^+^ T cells from 0.2% in NAT to 8% in the TME, while CD8^+^ effector T cells (CD8 Teff) constituted 36% of CD3^+^ T cells in NAT and 9% in the TME (**Figure 1c**). This relative increase in CD8 Texh was confirmed using a flow cytometry-based quantification of the CD39^+^PD1^+^ CD8 Texh population in the TME (**Extended Data Figure 1l and 2a**). We also identified Treg cells by high gene scores for Treg signature loci such as *IL2RA* and *FOXP3* (**Figure 1e**), as well as Treg cell-associated effector molecules such as *ENTPD1*, *CTLA4*, *GZMB*, and *IL-10* (**Extended Data Figure 2e**). CCR8^+^ tissue Treg cells were further sub-classified using the marker genes BATF and CCR8 (**Figure 1e**).

These findings demonstrate that single-cell chromatin accessibility landscapes can be used to decipher the immune cell composition in both tumor and normal tumor-adjacent tissue.

### Core- and tissue Treg signature characterize tissue Treg cells in tumor & NAT

Next, we performed the aforementioned analysis on two additional kidney tumor patients and annotated the immune landscape in TME and NAT (**Figure 2a** and **Extended Data Figure 2a-e)**. Due to experimental batch effects and high variations in the TME, we decided to analyze and display each patient individually. We then focused on Treg cells and used a chromatin-based “core Treg cell signature” that was originally computed by comparing Treg versus Tconv cell peak sets from human blood of healthy donors (data extracted from ^38^). We plotted this signature on all cell types identified in the tissue (**Figure 2b**, left, signatures listed in **Supplementary Table S1**). High core Treg epigenetic signature values were detected in CCR8^+^ Treg cells, but not in other CD4^+^ or CD8^+^ T cell populations, independently confirming our Treg cell type annotation. We then wondered how the core Treg signature was distributed in individual CCR8^+^ Treg cells of TME vs NAT. To address this, we ordered CCR8^+^ tissue Treg cells by signature rank and highlighted their tissue origin using a color code (**Figure 2c**). We computed the median rank position of tumor (red), NAT (grey), and the combined dataset (black) and observed no strong deviations from the combined dataset median. This suggests that the core Treg identity was unaffected by the inflammatory environment of the TME.

**Figure 2:**
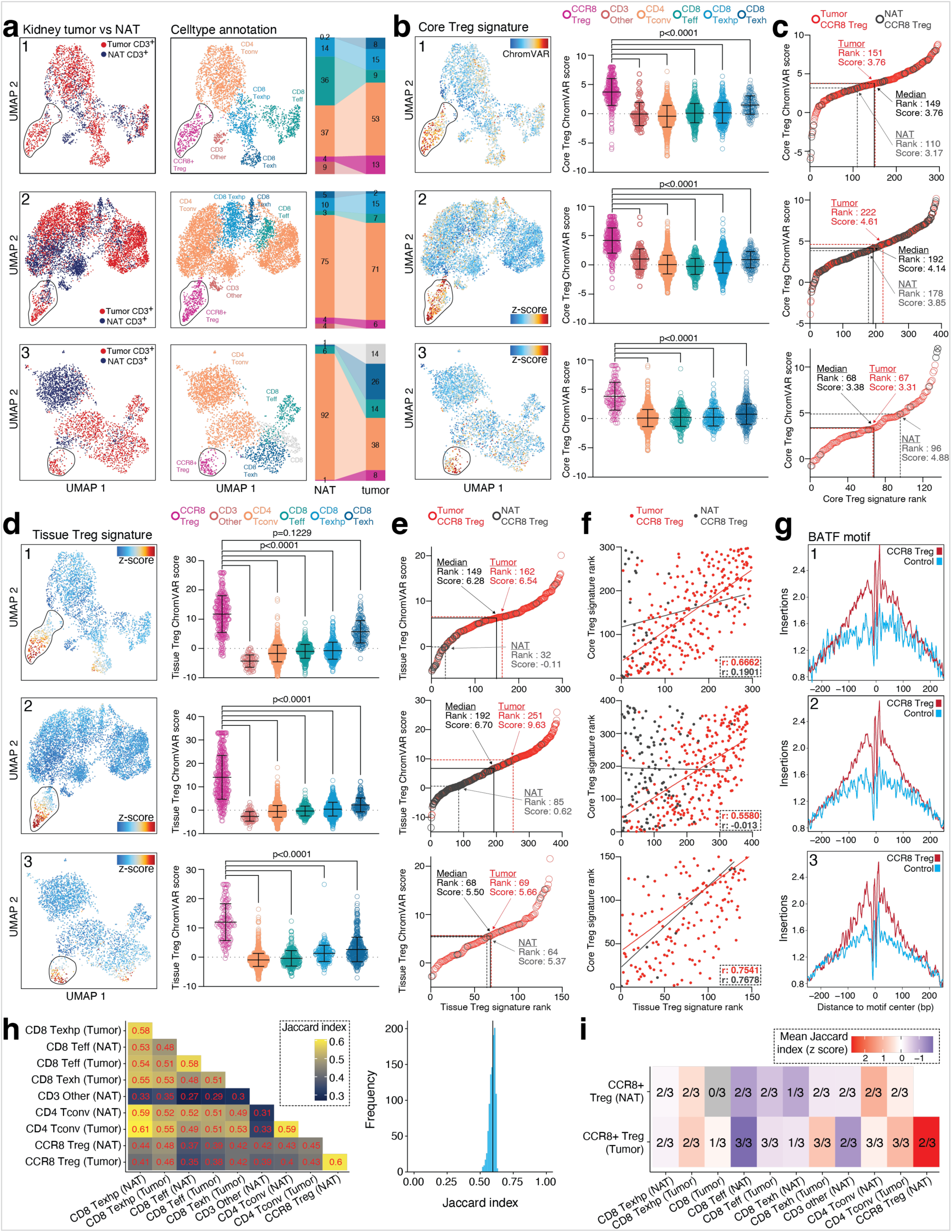
scATAC-seq of CD3^+^ T cells from kidney tumors. **(a)** UMAP and cell type annotation for three individual kidney tumor patients, with tumor and NAT CD3^+^ T cells. Patient 1 = clear cell renal carcinoma (CCRC), patient 2 = Renal angiomyolipoma (AML), patient 3 = CCRC. Left, origin of cell (tumor vs NAT); middle, celltype annotation; right, relative celltype composition presented as stacked bar graph. CCR8^+^ Treg cell cluster is circled. **(b)** Core Treg signature intensity for different cell types, Kruskal-Wallis and Dunn’s multiple comparisons test. **(c)** CCR8^+^ Treg cells were extracted and ordered by core Treg signature rank. Tumor and NAT CCR8^+^ Treg cells were then identified and highlighted in the dataset. The median rank for tumor (red), NAT (grey) and the combined dataset (black) was determined and the corresponding score was labeled in the figure. “x” highlights clipping on the y axis. **(d)** Tissue Treg signature intensity for different cell types, Kruskal-Wallis and Dunn’s multiple comparisons test. **(e)** CCR8^+^ Treg cells were extracted and ordered based on tissue Treg signature rank, as in (c). **(f)** Correlation between tissue Treg signature rank (x-axis) and core Treg signature rank (y-axis) for tumor CCR8^+^ Treg (red) and NAT CCR8^+^ Treg (black). Line indicates simple linear regression with corresponding Pearson correlation coefficient values shown in figure. **(g)** Computation of the BATF motif in the comparison of CCR8^+^ Treg cells versus a control population from the same dataset (e.g. CD4^+^ Tconv). **(h)** Left, Jaccard indices for patient 1. Only cell types with sufficient cell numbers for Jaccard index computation are included. Right, distribution of expected Jaccard indices assuming equality between tumor CCR8+ Treg cells and NAT CCR8+ Treg cells. Vertical lines mark the observed Jaccard index between these two populations. **(i)** Jaccard indices between accessible peaks in the different cell types (scaled mean across patients). Numbers indicate in how many of the three patients a Jaccard index could be computed. Gray fields indicate that a Jaccard index could not be computed in any patient. Additional flow cytometry data, scATAC-seq QC and cell type annotation in **Extended Data Figure 2**. Data representative of three independent experiments with three individual kidney tumor patients.

However, the question remains whether the TME affects the tissue program. Currently, no chromatin signatures for CCR8+ Treg cells of healthy liver or kidney donors exist. Therefore, we used a chromatin-based tissue regeneration signature derived from the comparison of CCR8^+^ tissue Treg cells isolated from tumor-free fat and skin tissue from healthy donors in comparison with CD45RA^+^ Treg cells isolated from blood (data extracted from ^35^). We then displayed the signature score for all tissue T cell types, as before, and observed high signature scores for CCR8^+^ tissue Treg cells in our dataset (**Figure 2d**). CCR8+ Treg cells from the tumor displayed higher overall tissue Treg signature scores than CCR8+ Treg cells from the NAT (**Figure 2e**). These findings suggest the possibility that, in an inflammatory TME, the tissue Treg program is more active than in the cognate NAT, potentially fueled by the presence of self-antigen. To investigate whether the core Treg signature, related to immune-regulatory activity, and the tissue Treg signature, related to the regenerative program, are correlated, we plotted core Treg signature rank vs tissue Treg signature rank for all three patients and observed positive Pearson correlation coefficients of r=0.5580, r=0.6662 and r=0.7551 for the TME (**Figure 2f**). Thus, we could hypothesize that the inflammatory environment in the TME fuels regenerative features of tissue Treg cells. For tissue Treg cells from fat and skin, it was shown that the regenerative program is linked to the transcription factor BATF in mouse^44, 45^ and human^35^. Therefore, we computed the impact of BATF on our dataset and observed a CCR8^+^ Treg cell specific BATF footprint in the epigenetic framework of those cells (**Figure 2g**).

So far, we have shown that CCR8^+^ Treg cells in the TME retain their core Treg epigenetic identity and show increased tissue Treg signature values. But are tumor CCR8^+^ Treg cells substantially different from CCR8^+^ Treg cells in the cognate NAT? To investigate this, we examined the overlap between accessible peaks in the analyzed cell types using Jaccard indices. For kidney tumor patient 1, among all comparisons involving CCR8+ Treg cells from tumor or NAT, the Jaccard index was highest between tumor CCR8+ Treg cells and NAT CCR8+ Treg cells (**Figure 2h**, left), and the observed Jaccard indices for these two populations were in the middle of the range of expected Jaccard indices under the assumption of cell type equality (**Figure 2h**, right histogram). For all other donors, Jaccard indices were computed (**Extended Data Figure 2f**) and summarized in a single comparison (**Figure 2i**).

These findings indicate that tumor Treg cells might not constitute a unique Treg cell type, but that the TME might harbor a strongly activated version of a tissue Treg cell.

### The core Treg epigenetic signature is not affected by the TME

In the previous paragraph, we identified a shared chromatin accessibility landscape of CCR8^+^ Treg cells in tumor vs cognate NAT. Besides chromatin features, Treg cells are also characterized by a Treg-specific DNA methylation pattern, which is considered a stable epigenetic mark of cellular memory^27, 39, 40^. Therefore, we generated tagmentation-based whole genome bisulfite sequencing data of CCR8^+^ Treg cells from TME versus NAT of three kidney tumor patients and analyzed 27,999,538 CpGs with a median coverage between 3 and 5 per sample (**Figure 3a** and **Extended Data Figure 3**). We extracted a core Treg DNA hypo- and hypermethylation signature, originally computed by comparing DNA methylation of CD45RA^+^CD127^-^CD25^+^ Treg versus CD45RA^+^CD127^+^CD25^-^ Tconv cells from human blood of healthy donors (data extracted from ^38^). This signature was then plotted on DNA methylation values of CCR8^+^ Treg cells isolated from tumor and NAT, and was displayed in a heatmap (**Figure 3b**, left heatmap). We also included pseudobulk replicates of single-cell chromatin accessibility data for these populations (**Figure 3b**, right heatmap). We then computed whether each region in tumor or NAT CCR8^+^ Treg matches the Treg hypomethylation pattern (2,242 regions), thus confirming Treg identity. In addition, we tested whether elements of the Tconv-specific hypomethylation pattern (1,419 regions) were methylated in Treg cells isolated from kidney tumor and NAT CCR8^+^ Treg cells (**Figure 3b**, bar graph). Our results indicate that between 80.0% and 93.8% of the loci from the Treg DNA hypomethylation pattern place tumor and NAT CCR8^+^ Treg cells closer to healthy blood Treg cells than to healthy blood Tconv cells. For the Tconv DNA hypomethylation pattern, this number ranged from 79.7% to 85.0%, confirming the epigenetic identity of our tumor and NAT cells. To show examples for core Treg- specific demethylated regions, we plotted the *FOXP3*, *CTLA4*, and *MIR146A*/*MIR3142* locus and overlaid methylation values for blood Treg, blood Tconv, tumor, and NAT CCR8^+^ Treg cells (**Figure 3c**). A direct comparison of the signature values between tumor CCR8^+^ Treg and cognate NAT CCR8^+^ Treg cells revealed no differences, indicating that the Treg-specific DNA methylation pattern was present in both TME and NAT.

**Figure 3:**
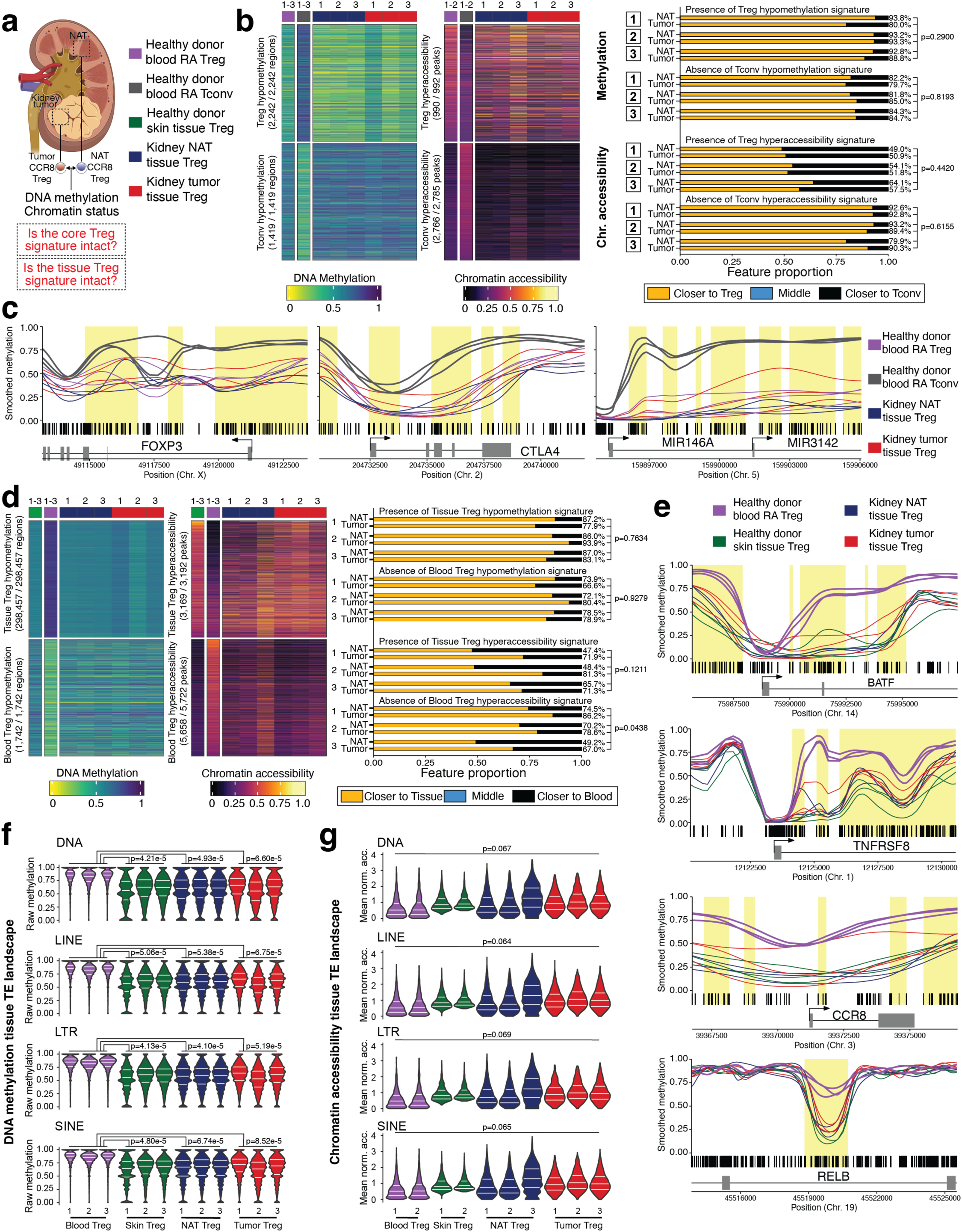
Whole-genome DNA methylation of liver tumor vs NAT CCR8^+^ Treg cells. **(a)** Overview of experimental design. **(b)** Left, heatmaps illustrating methylation (left) and chromatin accessibility (right, clipped at 95^th^ percentile) of features that are differential between blood CD45RA^+^ Tconv cells and blood CD45RA^+^ Treg cells. Columns indicate donors (Donors are aggregated for healthy blood cell types). Numbers indicate how many regions from the original signatures could be transferred to the present data set. Right, statistics related to heatmaps indicating whether tumor and NAT CCR8^+^ Treg cells are closer to blood CD45RA^+^ Treg cells or closer to blood CD45RA^+^ Tconv cells with respect to a feature. P values, two-tailed paired t test. **(c)** Smoothed methylation values for selected regions with blood CD45RA^+^ Tconv, blood CD45RA^+^ Treg, kidney tumor CCR8^+^ Treg and kidney NAT CCR8^+^ Treg. **(d)** Left, heatmaps illustrating methylation (left) and chromatin accessibility (right, clipped at 95^th^ percentile) of features that are differential between blood CD45RA^+^ Treg cells and skin tissue Treg cells. Columns indicate donors (Donors are aggregated for healthy-donor cell types). Numbers indicate how many regions from the original signatures could be transferred to the present data set. Right, statistics related to heatmaps indicating whether tumor or NAT CCR8^+^ Treg cells are closer to skin tissue Treg cells or closer to blood CD45RA^+^ Treg cells with respect to a feature. P values, two-tailed paied t test. **(e)** Smoothed methylation values for selected regions with blood CD45RA^+^ Treg, skin tissue Treg, kidney tumor CCR8^+^ Treg and kidney NAT CCR8^+^ Treg. **(f-g)** Methylation and chromatin accessibility of TE sites that belong to the four major TE classes and are hypomethylated or hyperaccessible in skin Treg cells from healthy donors. White horizontal lines indicate the lower and upper quartiles as well as the median. P values, One-way ANOVA with Tukey’s post-hoc test (Results of post-hoc test only shown in case of significant ANOVA). Data representative of independent experiments with individual kidney tumor patients (chromatin accessibility, n=3, DNA methylation, n=3).

For chromatin accessibility, we investigated both Treg hyper – and hypoaccessibility signatures. In this comparison, between 49.0% and 64.1% of the 990 peaks from the Treg hyperaccessibility signature placed tumor and NAT CCR8^+^ Treg cells closer to healthy blood Treg cells than to healthy blood Tconv cells. For the 2,766 peaks from the Tconv hyperaccessibility signature, this number ranged from 79.9% to 93.2%. Thus, our data indicate that the core Treg cell identity of tumor and NAT CCR8^+^ Treg cells on the chromatin accessibility level is pronounced but not as conserved as on the methylation level. Notably, we did not observe strong differences between tumor and NAT. Thus, we summarize that the core Treg-specific DNA methylation and chromatin accessibility patterns are similar between Treg cells from tumor as compared to NAT.

### CCR8^+^ Treg from tumor & NAT share tissue Treg epigenetic and TE landscapes

Next, we investigated the tissue Treg hypomethylation pattern consisting of 298,457 regions, initially computed by comparing tissue Treg cells isolated from healthy skin tissue in comparison with blood CD45RA^+^ Treg cells^38^, as well as the corresponding blood Treg hypomethylation pattern (1,742 regions). From the almost 300,000 regions that describe tissue Treg cells in healthy tissues, 77.9% to 93.9% (tissue Treg hypomethylation signature) and 72.1% to 80.4% (blood Treg hypomethylation signature) were intact in kidney CCR8^+^ Treg cells, with no differences between tumor tissue and cognate NAT (**Figure 3d**). In addition to the presence of the tissue Treg methylation signature, we also observed an overall trend towards hypomethylation across all chromosomes in tumor and NAT CCR8^+^ Treg cells, similar to what can be found in skin tissue Treg cells from healthy donors (**Extended Data Figure 3e**). To exemplify the high similarity between tissue Treg cells isolated from healthy donor skin, tumor, and NAT, we display selected regions for the tissue Treg DNA methylation pattern such as *BATF*, *TNFRSF8*, *CCR8*, or *RELB* (**Figure 3e**). On the chromatin accessibility level, we used a tissue Treg signature^38^ that not only contains genomic regions with hyperaccessibility in tissue Treg cells but also regions displaying hyperaccessibility in blood Treg cells. TME-derived tissue Treg cells showed a trend towards a stronger presence of the tissue Treg hyperaccessibility signature and a significantly stronger absence of the blood Treg hyperaccessibility signature, a finding consistent with tissue Treg signature ranks displayed before (**Figure 2e-f**).

Besides the aforementioned epigenetic tissue Treg signatures, we have previously shown that the epigenetics of TEs is another prominent feature characterizing tissue Treg cells^38^. Thus, we examined TE insertion sites that belonged to the four major TE classes (DNA, LINE, LTR and SINE) and had previously been shown to be demethylated in skin Treg cells compared to blood CD45RA^+^ Treg cells. (**Figure 3f**). An analogous analysis was performed on the chromatin accessibility level, focusing on TE insertion sites that are hyperaccessible in skin Treg cells. (**Figure 3g**). In these analyses, we found that both tumor and NAT CCR8^+^ Treg cells reflect the TE-specific epigenetic traits of healthy skin Treg cells although this reflection was only partly observed on the chromatin accessibility level. Notably, tumor and NAT CCR8^+^ Treg cells did not exhibit major differences in their TE-related epigenetic properties.

Together, these analyses indicate that DNA methylation, chromatin accessibility and the TE methylation landscape are highly similar between CCR8^+^ tissue Treg cells isolated from the tumor or cognate NAT, suggesting that both cell types might share a common ancestry.

### Similar chromatin accessibility in liver tumor and NAT CCR8^+^ Treg cells

In the previous experiments, we investigated tissue Treg cells in kidney tumors and the cognate NAT, and identified that, on chromatin accessibility-, DNA methylation- and TE level, these cell types appear very similar. A high similarity between tumor and NAT tissue Treg cells would have important implications in tumor Treg cell-targeting cancer therapies. We therefore set out to confirm our findings from the kidney in a different entity, namely primary hepatocellular carcinoma (HCC, **Figure 4** and **Extended Data Figure 4**). Comparable to the kidney, the cell type composition varies between tumor and NAT, with an increase of CCR8^+^ tissue Treg and CD8^+^ Texh cells in the tumor as compared to the NAT (**Figure 4a**), a finding confirmed by flow cytometry (**Extended Data Figure 4a**). Then, as before, we used the core- and tissue Treg signature to confirm the identity of the CCR8^+^ Treg population (**Figure 4b, d**) and ranked cells according to their signature score. We observed only minor differences between both tissues (**Figure 4c, e)**, and core- and tissue Treg signature ranks showed a positive correlation between 0.5044 and 0.5631 for the TME (**Figure 4f**). The BATF motif was enriched in CCR8^+^ Treg cells, highlighting the relevance of this factor – not only for tissue Treg cells in the kidney (**Figure 2g**), but also the liver (**Figure 4g**). The low total number of CCR8+ Treg cells in the NAT prevented computation of Jaccard indices between tumor CCR8+ Treg cells and NAT CCR8+ Treg cells, other Jaccard indices are shown in **Extended Data Figure 4f**.

**Figure 4:**
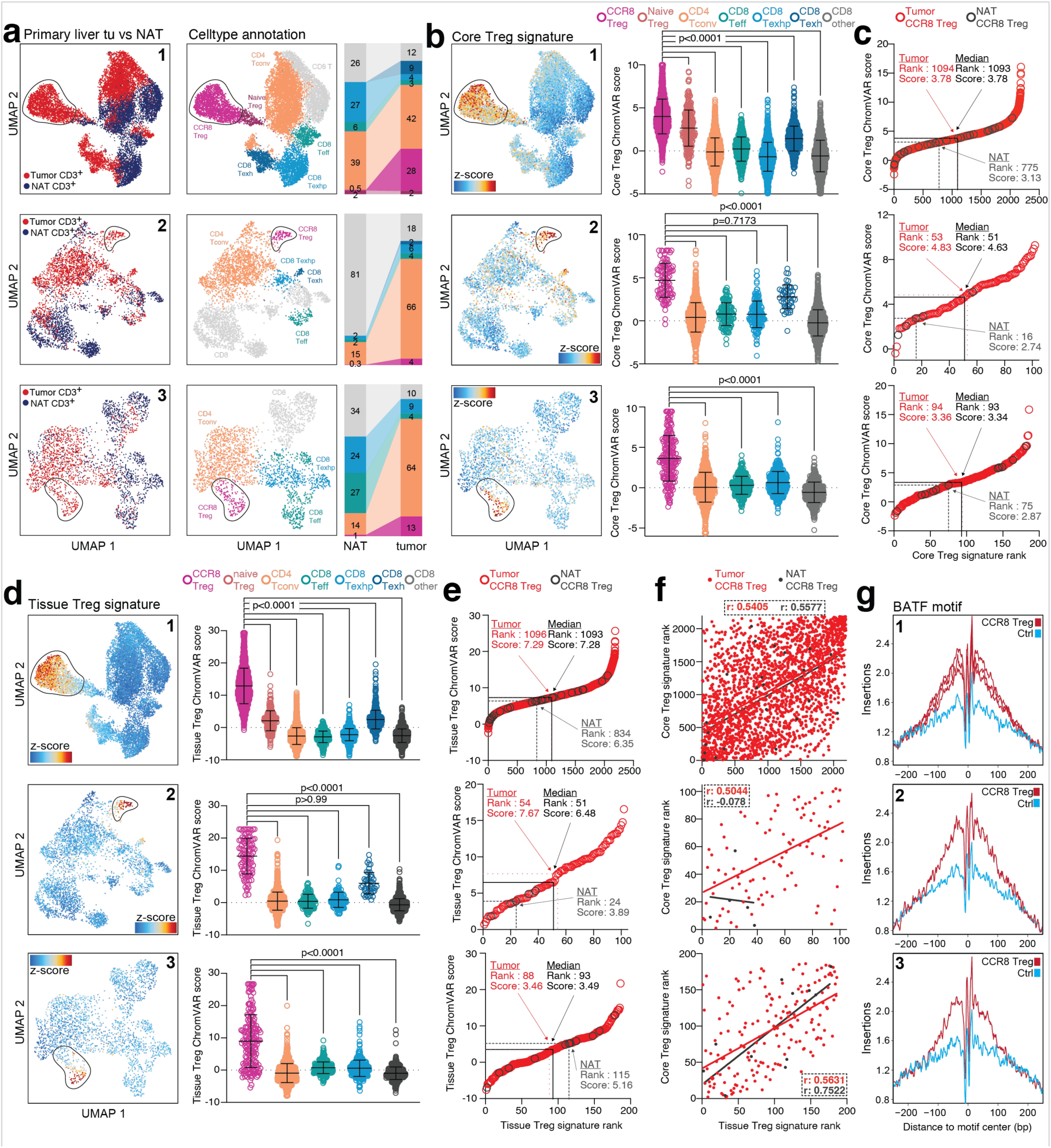
scATAC-seq of CD3^+^ T cells from liver tumors. **(a)** UMAP and celltype annotation for three individual liver tumor patients, with tumor and NAT CD3^+^ T cells. Patient 1-3: hepatocellular carcinoma. Left, origin of cell (tumor vs NAT); middle, celltype annotation; right, relative celltype composition presented as stacked bar graph. CCR8^+^ Treg cell cluster is circled. **(b)** Core Treg signature intensity for different cell types, Kruskal-Wallis and Dunn’s multiple comparisons test. **(c)** CCR8^+^ Treg cells were extracted and ordered based on core Treg signature rank. Tumor and NAT CCR8^+^ Treg cells were then identified and highlighted in the dataset. The median rank for tumor (red), NAT (grey) and the combined dataset (black) was determined and the corresponding score labeled in the figure. **(d)** Tissue Treg signature intensity for different cell types, Kruskal-Wallis and Dunn’s multiple comparisons test. **(e)** CCR8^+^ Treg cells were extracted and ordered based on tissue Treg signature rank, as in (c). **(f)** Correlation between tissue Treg signature rank (x-axis) and core Treg signature rank (y-axis) for tumor CCR8^+^ Treg (red) and NAT CCR8^+^ Treg (black). Line indicates simple linear regression with corresponding Pearson correlation coefficient values shown in figure. **(g)** Computation of the BATF motif in the comparison of CCR8^+^ Treg cells versus a control population from the same dataset (*e.g.* CD4^+^ Tconv).

Taken together, our data indicate that, also in primary hepatocellular carcinoma, CCR8^+^ Treg cells in tumor and NAT share a common epigenetic framework.

### DNA methylation & TE epigenetics suggest common ancestry of liver tumor & NAT CCR8^+^ Treg cells

Next, we generated tagmentation-based whole genome bisulfite sequencing data of CCR8^+^ Treg cells from TME versus NAT of 3 liver tumor patients, as before, and integrated these data with chromatin accessibility data. Our results indicate that the core Treg signatures were largely unaffected by the TME on both DNA methylation (**Figure 5a** and **Extended Data Figure 5a-d**) and chromatin accessibility level (**Figure 5b**), highlighted by the uniform DNA methylation patterns of *FOXP3*, *CTLA4*, or *MIR146A*/*MIR3142* (**Figure 5c**). Analogously, the tissue Treg signature was present in both liver tumor and NAT CCR8^+^ Treg cells without significant differences on both chromatin accessibility and DNA methylation level (**Figure 5d**). This similarity between CCR8^+^ Treg cells from liver tumor, liver NAT and skin tissue was observed also on a global scale (**Extended Data Figure 5e**), similar to our findings from kidney samples (**Extended Data Figure 3e**). The *BATF*, *TNFRSF8*, *CCR8*, and *RELB* gene methylation highlights the similarity between skin, liver NAT, and liver tumor CCR8^+^ Treg (**Figure 5e**). Finally, an investigation of the TE-related epigenetic traits reported in skin tissue Treg cells revealed no significant differences between tumor and NAT (**Figure 5f, g**).

**Figure 5:**
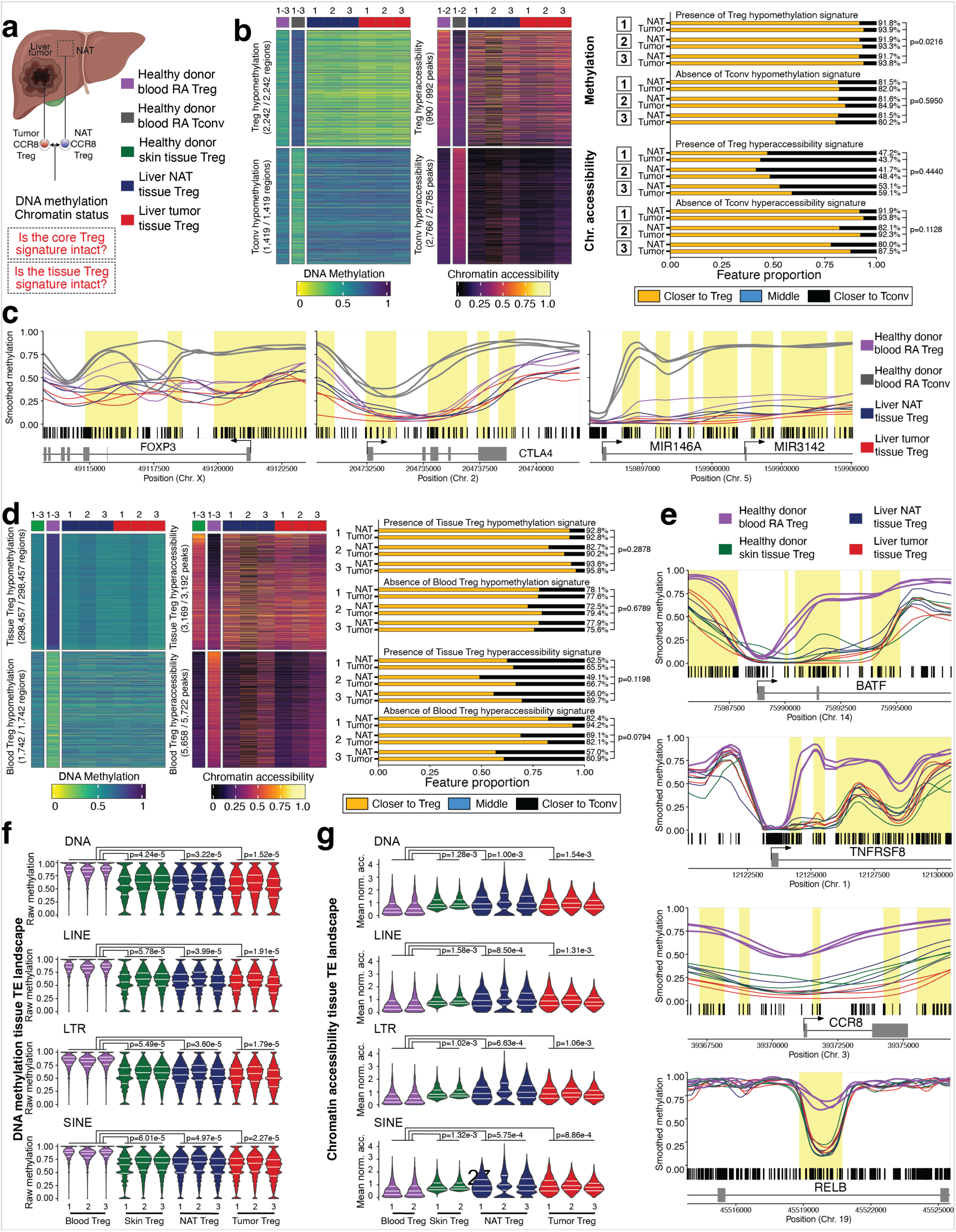
Whole-genome DNA methylation of liver tumor vs NAT CCR8^+^ Treg cells. **(a)** Overview of experimental design. **(b)** Left, heatmaps illustrating methylation (left) and chromatin accessibility (right, clipped at 95^th^ percentile) of features that are differential between blood CD45RA^+^ Tconv cells and blood CD45RA^+^ Treg cells. Columns indicate donors (Donors are aggregated for healthy blood cell types). Numbers indicate how many regions from the original signatures could be transferred to the present data set. Right, statistics related to heatmaps indicating whether tumor or NAT CCR8^+^ Treg cells are closer to blood CD45RA^+^ Treg cells or closer to blood CD45RA^+^ Tconv cells with respect to a feature. P values, two-tailed paired t test. **(c)** Smoothed methylation values for selected regions with blood CD45RA^+^ Tconv, blood CD45RA^+^ Treg, liver tumor CCR8^+^ Treg and liver NAT CCR8^+^ Treg. **(d)** Left, heatmaps illustrating methylation (left) and chromatin accessibility (right, clipped at 95^th^ percentile) of features that are differential between blood CD45RA^+^ Treg cells and skin tissue Treg cells. Columns indicate donors (Donors are aggregated for healthy-donor cell types). Numbers indicate how many regions from the original signatures could be transferred to the present data set. Right, statistics related to heatmaps indicating whether tumor or NAT CCR8^+^ Treg cells are closer to skin tissue Treg cells or closer to blood CD45RA^+^ Treg cells with respect to a feature. P values, two-tailed paired t test. **(e)** Smoothed methylation values for selected regions with blood CD45RA^+^ Treg, skin tissue Treg, liver tumor CCR8^+^ Treg and liver NAT CCR8^+^ Treg. **(f-g)** Methylation and chromatin accessibility of TE sites that belong to the four major TE classes and were previously reported to be differential between skin Treg cells and blood CD45RA^+^ Treg cells. White horizontal lines indicate the lower and upper quartiles as well as the median. Data representative of independent experiments with individual liver tumor patients (chromatin accessibility, n=3, DNA methylation, n=3).

Thus, these results indicate that DNA methylation, chromatin accessibility and the TE epigenetic landscape are highly similar also between CCR8^+^ tissue Treg cells isolated from liver tumors and the cognate NAT, further substantiating the hypothesis that both cell types share a common molecular framework.

### Tumor metastases harbor tissue Treg cells identical to target tissue

In the previous analysis, we identified shared molecular programs between tumor- and NAT-resident CCR8^+^ tissue Treg cells in primary kidney and primary liver tumor patients. However, since these primary tumors have developed in the same tissue that the NAT samples are extracted from, we cannot exclude that local inflammatory events already induce the tissue Treg phenotype even before a tumor can be clinically detected. This is especially the case for hepatocellular carcinoma, which is preceded by hepatitis (viral or non-viral) or cirrhosis^46^, but also kidney tumors can be associated with an inflamed tissue environment^47^. To disentangle the local tissue inflammation from our findings, we analyzed patients with metastases to the liver (**Figure 6** and **Extended Data Figure 6 and 7**). Since the malignancy did not arise in the liver but, in the case of our three patients, in the colon, the liver NAT is affected very little by the metastasis: the lobular structure of the liver parenchyma is maintained, there is a negligible degree of inflammatory infiltrate and no apparent steatosis (**Extended Data Figure 7a,** left). In contrast, the characteristic lobular structure is lost in the liver NAT of our HCC patients, and we can observe a portal as well intralobular inflammatory infiltrate and steatosis (**Extended Data Figure 7a,** right). The liver NAT from a metastasis patient thus is the closest we can get to a healthy corresponding tissue.

**Figure 6:**
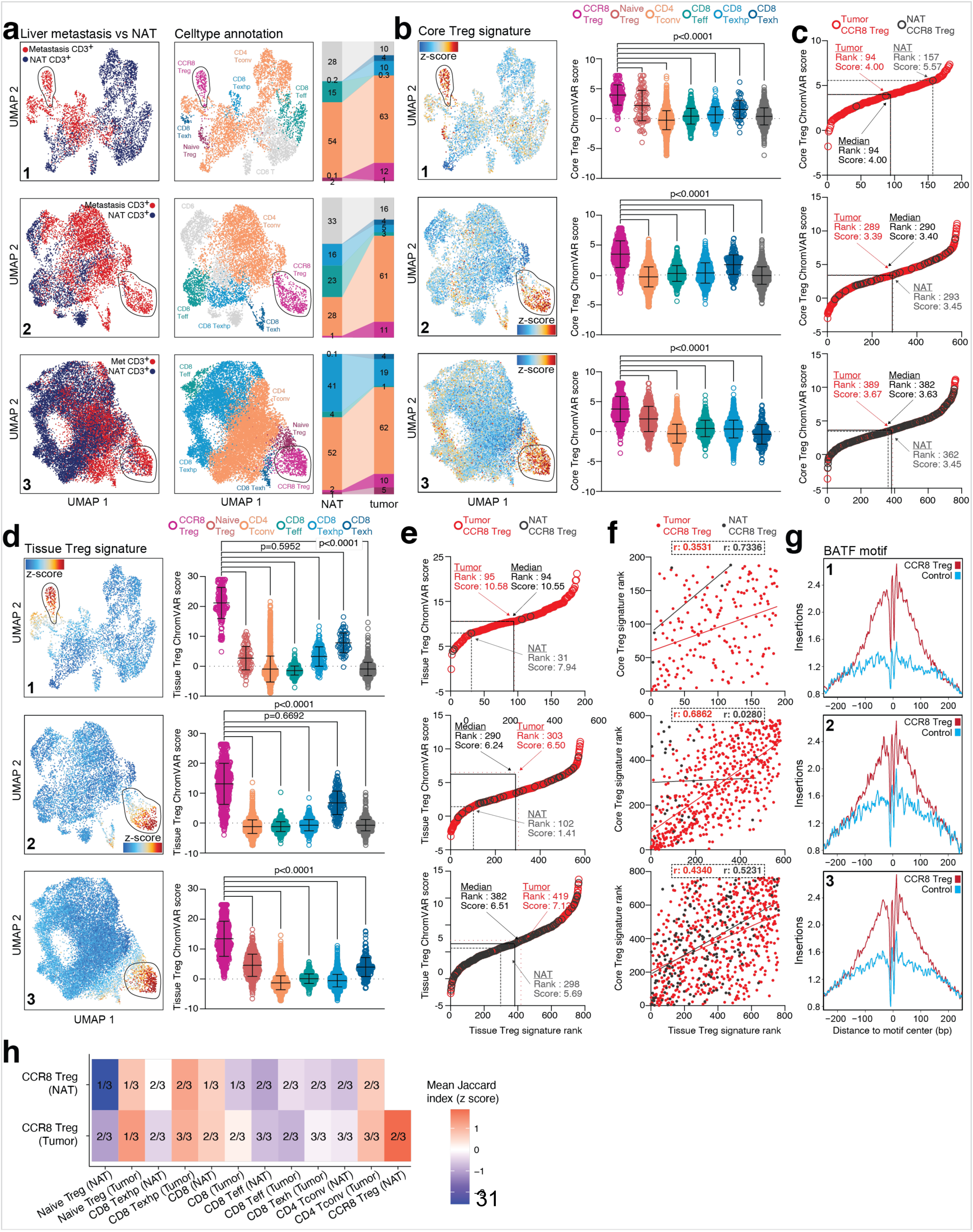
scATAC-seq of CD3^+^ T cells from metastasis to the liver. **(a)** UMAP and celltype annotation for three individual patients with metastasis to the liver, with tumor and NAT CD3^+^ T cells. Patient 1-3: metastasis to the liver from primary colorectal carcinoma. Left, origin of cell (tumor vs NAT); middle, celltype annotation; right, relative celltype composition presented as stacked bar graph. CCR8^+^ Treg cell cluster is circled. **(b)** Core Treg signature intensity for different cell types, Kruskal-Wallis and Dunn’s multiple comparisons test. **(c)** CCR8^+^ Treg cells were extracted and ordered based on core Treg signature rank. Tumor and NAT CCR8^+^ Treg cells were then identified and highlighted in the dataset. The median rank for tumor (red), NAT (grey) and the combined dataset (black) was determined and the corresponding score labeled in the figure. **(d)** Tissue Treg signature intensity for different cell types, Kruskal-Wallis and Dunn’s multiple comparisons test. **(e)** CCR8^+^ Treg cells were extracted and ordered based on tissue Treg signature rank, as in (c). **(f)** Correlation between tissue Treg signature rank (x-axis) and core Treg signature rank (y-axis) for tumor CCR8^+^ Treg (red) and NAT CCR8^+^ Treg (black). Line indicates simple linear regression with corresponding Pearson correlation coefficient values shown in figure. **(g)** Computation of the BATF motif in the comparison of CCR8^+^ Treg cells versus a control dataset from the same dataset (*e.g.* CD4^+^ Tconv). **(h)** Jaccard indices between accessible peaks in the different cell types (scaled mean across patients). Numbers indicate in how many of the three patients a Jaccard index could be computed. Gray fields indicate that a Jaccard index could not be computed in any patient. Data representative of three independent experiments with three individual liver metastasis patients.

Comparable to the kidney and primary liver tumor samples, we also observe a difference in cell type composition between the metastasis and the corresponding liver NAT in the three liver metastasis patients (**Figure 6a**) and confirmed these findings via flow cytometry (**Extended Data Figure 6b**). The frequency of CCR8^+^ tissue Treg cells is increased in the metastasis compared to the NAT, as is the frequency of CD8 Texh. We can further observe a decrease of CD8 Teff cells in the metastasis compared to the NAT. CCR8^+^ tissue Treg cell clusters from all three patients are characterized by high z-scores for the core Treg cell signature (**Figure 6b**) and the tissue Treg cell signature (**Figure 6d**). Then, as before, we ranked cells according to their signature score, which showed only minor differences between signature values for tumor and NAT (**Figure 6c, e**), and core and tissue Treg signature ranks showed a positive correlation between 0.3531 and 0.6862 for tumor and 0.0280 and 0.7336 for NAT (**Figure 6f**). We observed a strong BATF footprint in the CCR8^+^ Treg cells cluster compared to a control population (*e.g.* CD4^+^ Tconv) (**Figure 6g**). Jaccard indices of accessible peaks were highest between tumor CCR8^+^ Treg cells and NAT CCR8^+^ Treg cells (**Figure 5h** and **Extended Data Figure 5f**).

Taken together, these results suggest common molecular characteristics in CCR8^+^ Treg cells in tumor metastasis compared to healthy tissue even without accompanying tissue inflammation.

### Tissue Treg cells share characteristics across primary tumor and metastasis

So far, our findings have strongly indicated that CCR8^+^ Treg cells in tumor and the cognate NAT are governed by the same regulatory program. Therefore, we now wanted to analyze whether there are any differences in the chromatin accessibility of CCR8^+^ Treg cells between the primary tumor and the corresponding metastasis of the same individual. To investigate this, we processed a colorectal carcinoma sample with the corresponding colon NAT along with the metastasis to the liver and the corresponding liver NAT, all from one patient (**Figure 7a** and **Extended Data Figure 7**). The cell type composition differs between the tumor and colon NAT, as well as between the liver metastasis and liver NAT, with a higher frequency of CCR8^+^ tissue Treg cells in malignant tissue as compared to the NAT (**Figure 7b**), confirmed via flow cytometry (**Extended Data Figure a-b**). Two CCR8^+^ Treg cell clusters were identified, one containing cells mainly from the CRC, the other containing cells mainly from the liver metastasis. Both clusters have high z-scores for the core Treg cell signature as well as the CCR8^+^ Treg cell signature (**Figure 7c, e**), and signature ranks indicate that both signatures are present more strongly in malignant tissue compared to the NAT (**Figure 7d, f**). The positive correlation can be observed in primary tumor and metastasis, irrespective of the anatomical location (**Figure 7g**).

**Figure 7:**
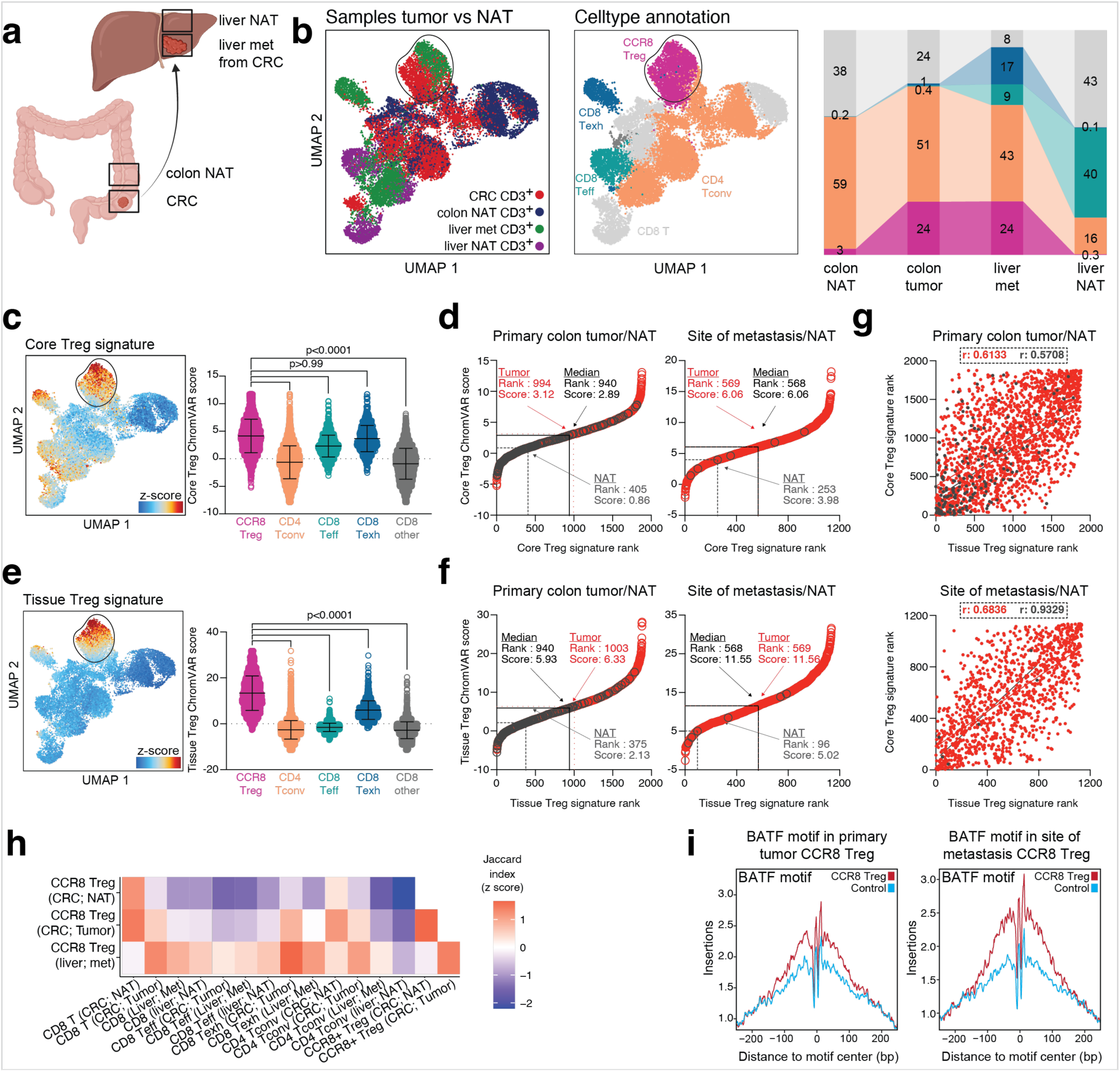
scATAC-seq of CD3^+^ T cells from primary tumor and metastasis. **(a)** Experimental overview. **(b)** UMAP and cell type annotation. Left, origin of cell (tumor vs NAT, metastasis vs primary tumor); middle, celltype annotation; right, relative celltype composition presented as stacked bar graph. CCR8^+^ Treg cell cluster is circled. **(c)** Core Treg signature intensity for different cell types, Kruskal-Wallis and Dunn’s multiple comparisons test. **(d)** CCR8^+^ Treg cells were extracted and ordered based on core Treg signature rank for primary tumor or site of metastasis. Tumor and NAT CCR8^+^ Treg cells were then identified and highlighted in the dataset. The median rank for tumor (red), NAT (grey) and the combined dataset (black) was determined and the corresponding score labeled in the figure. **(e)** Tissue Treg signature intensity for different cell types, Kruskal-Wallis and Dunn’s multiple comparisons test. **(f)** CCR8^+^ Treg cells were extracted and ordered based on tissue Treg signature rank, as in (d). **(g)** Correlation between tissue Treg signature rank (x-axis) and core Treg signature rank (y-axis) for tumor CCR8^+^ Treg (red) and NAT CCR8^+^ Treg (black). Line indicates simple linear regression with corresponding Pearson correlation coefficient values shown in figure. **(h)** Jaccard indices (scaled) between accessible peaks in the analyzed cell types. Grey fields indicate that a Jaccard index could not be computed due to low cell numbers. **(i)** Computation of the BATF motif in the comparison of colon CCR8^+^ Treg cells versus a control population from colon (*e.g.* CD4^+^ Tconv) (left) and liver CCR8^+^ Treg cells versus a control population from the liver (*e.g.* CD4^+^ Tconv) (right). Data representative of a single experiment with one individual tumor patient.

When we analyzed cell type similarity measured by Jaccard indices, we found that the similarity between tumor and NAT CCR8^+^ Treg cells as well as the similarity between Tumor CCR8^+^ Treg cells from primary tumor and metastasis was among the highest observed similarities invoving CCR8^+^ Treg cells (**Figure 7h**). The corresponding Jaccard indices were at the lower end of the range of expected indices assuming cell type equality (**Extended Data Figure 7h**). Chromatin footprints again show that, both in primary tumors and metastasis, BATF is a key TF in establishing the CCR8^+^ Treg epigenetic landscape (**Figure 7i**).

Thus, in summary, in primary tumors and metastasis only slight differences in chromatin accessibility between CCR8^+^ Treg cells were observed, indicating that core and tissue Treg chromatin accessibility might be independent from the anatomical location and shared between different tissues, promoting not only the idea of a common ancestry, but to a large extent even an universal tissue Treg signature^2^.

### The tissue Treg program promotes tumor spheroid growth *in vitro*

A detailed molecular analysis of chromatin accessibility, DNA methylation, and TE landscape revealed that CCR8^+^ Treg cells in tumors are very similar to CCR8^+^ Treg cells from the cognate tumor-free tissue. Irrespective of their tissue of origin (kidney, liver or colon) or the tissue microenvironment (TME vs healthy tissue), they carry a phenotypic imprint of the transcription factor BATF, a factor closely linked to the tissue Treg program. Given the BATF-supported tissue-remodeling effector function, we asked whether the tissue Treg program would also promote tumor spheroid growth. Due to the low overall numbers and the difficulty to culture TME-derived tissue Treg cells, we decided to induce BATF expression in blood-derived naive Treg cells for 6 days *in vitro* using a cytokine cocktail with IL-2, TGF-β, IL-12, IL-21, IL-23 and TCR stimulation using TransACT^35^. As control, we used expanded Treg cells treated only with IL-2 and TransACT. After six days, cells were washed vigorously to remove all traces of the cytokine mix, incubated overnight in fresh media, and cell-free supernatant was harvested. Intracellular flow cytometry revealed that BATF was specifically induced in tissue-like Treg cells treated with the cytokine cocktail, while FOXP3 expression levels remained high in both tissue-like and control Treg cell cultures (**Extended Data Figure Figure 8a**). Cell-free supernatant from both tissue-like and control Treg cells was added to tumor spheroid cultures in a 1:16 dilution, and spheroids were imaged hourly for several days. Although all spheroids were roughly equally sized one hour after experimental start (**Figure 8a**, left), tumor spheroids incubated with tissue-like Treg supernatant had increased to 1.012x10^3^ µm^2^ mean largest object area, while spheroids with control Treg supernatant grew 354x10^3^ µm^2^ mean largest object area (**Figure 8a**, right). Using both tissue-like Treg and control Treg supernatants at 1:64 dilution abrogated the effects (**Extended Data Figure 8b**), and incubation with the tissue Treg induction medium consisting of IL-2, TGF-β, IL-12, IL-21, and IL-23 mediated no spheroid growth-promoting effects (**Extended Data Figure 8c**). These results indicate that inducing a tissue-like Treg program promotes the secretion of factors that promote tumor spheroid growth.

**Figure 8:**
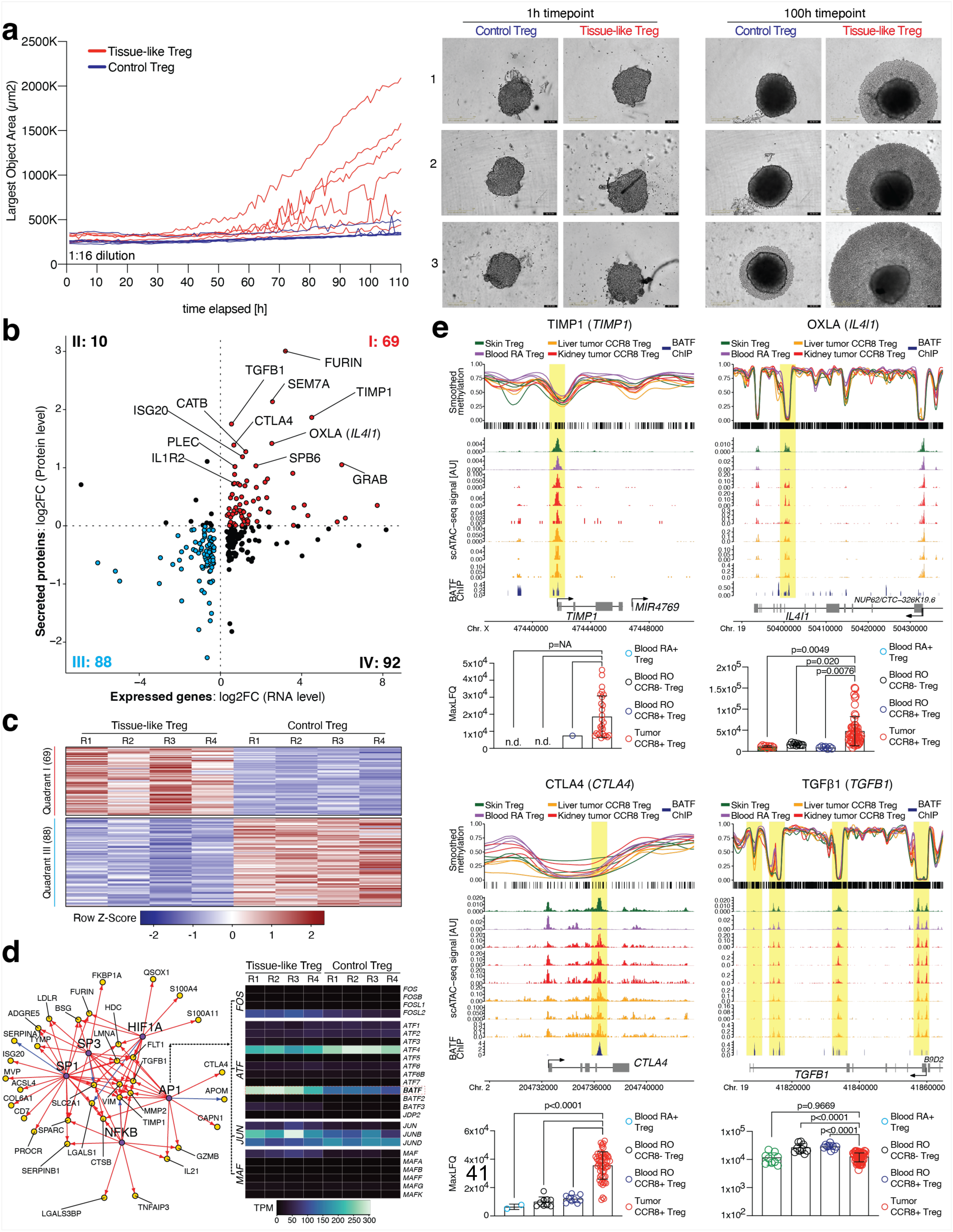
Inducing the tissue Treg program promotes tumor spheroid growth *in vitro*. **(a)** Human blood CD45RA^+^ Treg cells were treated for 6 days *in vitro* using TransAct, IL-2, TGF-β, IL-12, IL-21, IL-23 (Tissue-like Treg) versus TransAct and IL-2 (control Treg). Then, tumor spheroids were treated with cell culture supernatant at 1:16 dilution and imaged for 110 hours using an Incucyte imaging system. Representative images from 3 donors to the right (n=3 with two technical replicates). Additional controls **Extended Data Figure 8a-c**. **(b)** Correlation of gene expression data of tissue-like versus control Treg cells with quantitative proteomics of cell culture supernatant produced as shown in (a). For plotting, only significantly differential genes (n=4, Deseq2) have been correlated with the corresponding protein. Data are mean values derived from 4 donors (RNA) and 5 donors with three technical replicates each (secretome). **(c)** Heatmap highlighting the expression of genes from quadrant I (69) and quadrant III (88) in 4 individual replicates of tissue-like Treg and control Treg cells, row-scaled. **(d)** Left, gene regulatory network restricted to regulatory interactions with target genes that are up-regulated in tissue-like Treg cells both on the RNA level and on the protein level (quadrant I, 69 genes, the five transcription factors with the largest degree of connectedness are shown). Purple nodes represent transcription factors, yellow nodes represent transcription factor targets, red edges denote positive regulation, blue edges denote negative regulation. Right, expression of AP1 family members *FOS*, *ATF*, *JUN* and *MAF* in tissue-like Treg versus control Treg cells, global expression values. Data representative of three or more independent experiments with three or more individual donors or tumor patients. TPM, transcripts per million. **(e)** Smoothed methylation values for selected regions with blood CD45RA^+^ Treg, skin Treg, kidney tumor CCR8^+^ Treg and liver tumor CCR8^+^ Treg cells. Below, chromatin accessibility and BATF ChIP-seq signal in human GM12878 cells^51^. Lower end of each graph: results from quantitative proteomics of FACS-sorted blood CD45RA^+^ Treg, blood CD45RO^+^CCR8^-^ Treg, blood CD45RO^+^CCR8^+^ Treg and tumor CCR8^+^ Treg cells (n=3-19 with three technical replicates, one-way ANOVA).

### Quantitative proteomics reveals tissue and tumor Treg-associated factors

We next sought to identify the factors in the tissue Treg supernatant that could potentially promote tumor growth. We again generated tissue Treg versus control Treg supernatant and measured secreted proteins in the supernatant using quantitative proteomics. Proteome data were correlated with gene expression profiles of tissue-like Treg versus control Treg cells^35^ and displayed in a log2 fold change (FC) - log2FC plot (**Figure 8b** and **Supplementary Table S2**). In this comparison, 69 factors were identified for which gene expression was significantly upregulated and secreted proteins exhibited higher levels, while 88 genes showed reduced secretion (**Figure 8b-c**). Next, we used this list of 69 secreted factors (**Figure 8b**) and generated a directed graph of regulatory interactions, in which each transcription factor (TF) was linked to its target gene (**Figure 8d**, left). This analysis identified 5 central TFs; one factor, NFκB, is well known for its essential function in Treg cell development and function^48^; another factor, AP1, summarizes a group of several potent TFs of the *FOS*, *ATF*, *JUN* and *MAF* families^49^ and was directly linked to secreted molecules like CTLA4, TIMP-1 or TGF-β. To identify an AP1 family member that is likely responsible for the observed changes, we used transcriptomics data of tissue-like Treg cells and observed high expression values for *ATF4* and *BATF* TFs (**Figure 8d**, right). In the comparison of tissue-like Treg with control Treg cells, of those two TFs, only *BATF* was up-regulated upon induction of the tissue Treg program *in vitro*. Since BATF was also identified as a major TF shaping the tissue and tumor CCR8^+^ Treg epigentic landscape (**Figure 2-7**), we hypothesize that this factor might play an important role in promoting the tissue Treg tissue-remodeling and immunosuppressive effector program in the TME.

Among these molecules identified in our combined transcriptome/secretome analysis were proteins related to matrix remodeling, such as TIMP1, a matrix metalloproteinase associated with metastasis and the invasion potential of tumors^50^. To further investigate the relevance of those factors for the TME, we plot the DNA methylation and chromatin accessibility for selected candidates, and include quantitative proteomics data from FACS-sorted CCR8^+^ Treg cells from human tumors. This analysis highlights that *TIMP1* has conserved methylation and chromatin accessibility traits in healthy tissue Treg, liver and kidney tumor Treg samples, and that TIMP1 is produced in tumor-derived CCR8^+^ Treg cells compared to blood-based control Treg cells, where levels were below the detection limit (**Figure 8e**, top left). In addition, we extracted BATF ChIP-seq data^51^ and overlaid the BATF binding pattern, and observed a correlation between DNA demethylation, increased chromatin accessibility and BATF binding in the *TIMP1* promoter region.

Another factor identified in our combined transcriptome/secretome screen was OXLA (encoded by *IL4I1*), an aryl hydrocarbon receptor activating enzyme that promotes cancer cell motility and also suppresses adaptive immunity, thus playing a dual role in immunosuppression and cancer progression^52^. Also, this factor showed conserved epigenetic patterns, BATF binding in the gene body and increased expression in CCR8^+^ Treg cells isolated from tumors (**Figure 8e** top right). Besides TIMP1 and OXLA, we also identified other factors related to immune suppression, such as CTLA4 in its soluble form, which is involved in attenuating T cell activation, inhibiting immune cell differentiation, and modulating anti-tumor immunity^53, 54^; and TGF-β, a potent promotor of tumor progression, invasion, and metastasis^55^ (**Figure 8e**, bottom left and right). Other proteins idenfied in our screen include SEM7A, a membrane-anchored protein shown to promote growth, motility, invasion, as well as lymphangiogenesis in the TME of ductal carcinoma^56^; FURIN, a proprotein convertase essential for the maintenance of peripheral immune tolerance^57^; and IL1R2, a receptor that can be membrane-bound or secreted and that was described on Treg cells in the immunosuppressive TME of human cancer^58^ (**Extended Data Figure 8c**).

In summary, our combined secretome/transcriptome screen identified several candidate proteins involved in Treg-mediated immune suppression, tissue regeneration and cancer progression, many of which were related to the activity of AP1 transcription factors such as BATF. These candidate factors could constitute a reservoir of druggable targets for therapeutic interventions to combat both immunosuppression and tissue regeneration in the TME.

## DISCUSSION

Treg cells in non-lymphoid tissues have been found to promote tissue regeneration and homeostasis, an important non-canonical function in addition to their “classical” role in immune regulation. In the tumor microenvironment, Treg cells are generally considered detrimental: Meta-analysis of malignancies across a variety of tissues showed that a high level of Treg cell infiltration into the tumor correlates with decreased overall patient survival in most of the cancer types investigated, including melanoma, HCC, cervical, mammary, renal, and gastric cancers^47^. Also in this study, conversely, elevated Treg cell infiltration correlated with better outcomes in colorectal, head and neck, and esophageal cancers, pointing towards the need to better understand Treg cell heterogeneity in different tumors. Looking specifically at CCR8^+^ Treg cells, a high CCR8^+^ Treg cell frequency of total Treg cells was correlated with a poor prognosis in a breast cancer study^32^, as was a high expression of CCR8 in CRC and NSCLC samples, where tumor-infiltrating CCR8^+^ Treg cells were described as highly immunosuppressive^59^. In another study on NSCLC, single-cell profiling of Treg cells identified an IRF4-expressing population of tumor-infiltrating, highly suppressive Treg cells, reminiscent of CCR8^+^ Treg cells, and the presence of this cell type was further described to correlate with a poor prognosis in various human cancer types^60^. Retrospective analysis of samples from more than 250 patients with muscle-invasive bladder cancer (MIBC) explored the prognostic value of CCR8^+^ Treg cell levels on responsiveness to chemotherapy and overall survival. They found that the abundance of CCR8^+^ Treg cells in MIBC correlated inversely with responsiveness and overall survival^31^.

Based on these findings, therapies targeting “tumor Treg” cells are being developed, with promising results: CCR8-specific monoclonal antibodies designed to deplete CCR8^+^ Treg cells induce CD8 Teff cell infiltration into the tumor and lead to pro-inflammatory responses in a variety of tumor mouse models^28, 33, 61^. Combination of CCR8^+^ Treg cell targeting antibodies with anti-PD1 therapy has been shown to potentiate the anti-tumor effect in murine tumor models^30, 33, 61^. However, some degree of autoimmune inflammation has been observed upon CCR8^+^ Treg cell depletion, albeit at lower levels than by complete Treg cell depletion^33^.

In the study presented here, we performed a detailed characterization of Treg cells isolated from human tumors and the corresponding NAT. We found that CCR8^+^ Treg cells are present not only in the tumor, but also in the cognate NAT, albeit at lower frequencies. In order to assess molecular programs active in CCR8^+^ Treg cells isolated from tumor and NAT, we performed scATAC-seq and whole-genome bisulfite sequencing, and found that CCR8^+^ Treg cells from both tumor and healthy tissue shared epigenetic patterns including the core Treg program and the tissue regenerative program. In addition, the tissue-specific TE landscape including DNA, LINE, LTR, and SINE elements revealed no significant differences between tumor and NAT CCR8^+^ Treg cells. This suggests that “tumor Treg” cells are not a tumor-specific cell type, but that CCR8^+^ Treg cells found in tumor and healthy tissue are the same cell type, potentially with the same tissue-regenerative properties on top of the classical suppressive Treg cell program. To our knowledge, there is no comprehensive study that analyzed CCR8^+^ Treg cells in the tumor vs CCR8^+^ Treg cells from the corresponding healthy tissue using a combination of chromatin accessibility, DNA methylation and TE analysis.

A study on human lung cancer identified BATF as a key transcription factor for Treg cells in the TME comparing Treg cells isolated from the tumor to peripheral Treg cells isolated from blood^29^, which aligns with our data. CCR8^+^ Treg cells with similar transcriptional programs were also found in inflamed tissue from arthritic joint inflammation patients in a different study^62^, which agrees with our finding that CCR8^+^ Treg cells from inflamed NAT in HCC patients exhibit the BATF-dependent tissue-remodeling program. Also, coming back to data from murine studies, a landmark study systematically characterized murine Treg cell populations across lymphoid and non-lymphoid tissues using scRNA/TCR-seq, and found that Treg cell subsets were shared between many tissues, including the tissue Treg cell population being shared (and being transcriptionally comparable) especially between non-lymphoid tissues^2^. Parabiosis experiments revealed that CCR8^+^ Treg cells frequently leave and re-enter non-lymphoid tissues in a non-tissue-specific manner, with dwell times of about 3 weeks for tissue Treg cells in most non-lymphoid tissues^2^. Together, these findings suggest a dynamic model of tissue residency of CCR8^+^ Treg cells with multi-tissue homing in mice. Our data from human tumor and cognate NAT would fit well into this model, however further research is required to investigate CCR8^+^ Treg cell dynamics in human.

Since our results indicate that CCR8^+^ tissue Treg cells found in tumors have the same non-lymphoid tissue phenotype as the cognate tumor-free tissue, with intact core- and tissue Treg effector signatures on both chromatin accessibility- and DNA methylation level, directly targeting the CCR8 surface molecule therapeutically could also delete these populations in healthy tissues^38^. An alternative to directly targeting CCR8 on Treg cells could be to interfere with their selective recruitment to the tumor. If the dynamics observed in the murine system, namely a relatively short dwell time in non-lymphoid tissues of about 3 weeks, holds true for the human system, this might be a promising strategy.

In this study, we additionally performed quantitative proteomics of supernatant derived from *in-vitro* generated tissue-like Treg cells. This supernatant promotes tissue regeneration^35^, and using *in-vitro* tumor spheroid cultures we could show that the supernatant also promotes tumor growth. In our screen, we identified candidate factors linked to both the regenerative and suppressive program of tissue Treg cells, and confirmed the expression of factors such as *TIMP1*, a matrix metalloproteinase associated with tumor metastasis and invasion^50^, or *IL4l1*, an AHR activating enzyme promoting cancer cell motility and suppressing adaptive immunity^52^ also in primary human tumor-resident CCR8^+^ Treg cells. We could link the activity of those factors to 5 TF families including NFκB and AP1, which process signals also from the T-cell receptor. Thus, the autoreactive nature of (tissue) Treg cells could potentially be linked to their effector program, which includes not only immunosuppressive molecules, but also tissue-regenerative and tumor growth-promoting factors. A selective targeting of these tissue Treg effector molecules could constitute a therapeutical intervention strategy to combat tissue Treg programs in the TME without the systemic elimination of all effector Treg cells.

## ACKNOWLEDGEMENTS

Tissue samples were provided by the tissue bank of the University Medical Center Mainz in accordance with the regulations of the tissue biobank and the approval of the ethics committee of University Medical Center Mainz.

We acknowledge the DKFZ’s ODCF and GPCF core facilities for supporting the genomic sequencing and data processing. The authors gratefully acknowledge the computing time granted on the supercomputer MOGON NHR at Johannes Gutenberg University Mainz as part of NHR South-West (<u>nhrsw.de</u>). We would like to thank the Core Facility for Next-Generation Sequencing (CFNGS) at the FZI for their invaluable support. We would like to thank the FACS Core Facility CFFC PKZI FZI of the University Medical Center of the Johannes Gutenberg-University Mainz for providing support and instrumentation FACSAria III and FACSymphony A5 funded by the Deutsche Forschungsgemeinschaft (DFG, German Research Foundation) – INST 371/44-1 FUGB and INST 371/45-1 FUGB.

This work was funded by the Deutsche Forschungsgemeinschaft (DFG, German Research Foundation) Projektnummer 318346496 – SFB1292/2 TP19N (to M.D. and F.M.), TP11 (to U.D.) TP22N (to M.M.G.), TPQ1 (to S.T., M.M.G.), TP01 (to T.B.), and Projektnummer 490846870 – TRR355/1 TPA01 (to M.D.), TPA09 (to T.B.), TPA10 (to T.B.) and Projektnummer 446605368 - DFG priority program SPP 2225 (to U.D.).

## AUTHOR CONTRIBUTIONS

Study design: M.D.; Generation of main sequencing datasets: K.L.B. and T.K.; Analysis of scATAC-seq data: K.L.B. and N.B.; Analysis of bisulfite sequencing data: N.B.; Analysis of TEs: N.B; Contribution to experimental work and/or analyses: K.L.B., T.K., N.B., M.S., S.S.H., D.M.M., M.M.V., A.S.N., K.B., J.-P.M., U.D., S.T., M.V., D.W., C.P., M.B., M.L., S.D., M.K., T.B., H.-J.S., A.H., E.F., F.M., B.B., M.M.G.; Original manuscript draft: K.B., T.K., N.B., and M.D.; Supervision of investigation: M.D.; Funding acquisition: M.D.

## METHODS

### Human tissue

Human tissue samples were obtained from patients undergoing tumor resection at the University Medical Center Mainz. Tissue was characterized as tumor tissue or NAT by a pathologist, and, if possible, NAT samples were collected with spatial distance to the tumor tissue, since tissue in the vicinity of tumors is often inflamed or fibrotic. Experiments were approved by the Landesärztekammer Rheinland-Pfalz, ethics vote 2021-15834 (“Immunzelllandschaft im Gewebe”). Details on age, gender, diagnosis, and pathological characterization are listed below. Patients did not receive chemo-, radiation-, or immunotherapy prior to resection. For all samples provided in this manuscript, we will provide basic information about tumor type, age of the patient, gender, and specific pathological classification with **TNM** and **GLVPn** status, if applicable (**T**, tumor, grade 1-4 relates to size and degree of tissue infiltration; **N**, regionary lymph nodes, grade 0 = no lymph nodes affected, grade 1-3 relates to degree of tumor cells infiltrating lymph nodes; **M**, distant organ metastasis, grade 0 = no metastases found, grade 1 = metastases found; **G**, grade of differentiation, G1-4, degree of differentiation is anti-proportional, malignancy is proportional to the value of the number; **L**, lymph vessels affected (0/1); **V**, blood vessels affected (0/1); **Pn**, perineural tumor invasion (0/1)). All histological diagnoses were confirmed by board certified pathologists in the routine diagnostics of the Institute of Pathology at the University Medical Center Mainz.

Kidney tumor patients for single-cell chromatin accessibility sequencing (**Figure 1+2**) were classified as: (1) Clear cell renal carcinoma, 64 years, male, T1aN0M0-G2L0V0Pn0; (2) Renal angiomyolipoma, 62 years, female; (3) Clear cell renal carcinoma, 71 years, female, T3aN0M0-G2L0V1Pn0.

Kidney tumor patients for tagmentation-based whole genome bisulfite sequencing (**Figure 3**) were classified as: (1) Clear cell renal carcinoma, 44 years, female, T3aN0M0-G4L0V0Pn0; (2) Clear cell renal carcinoma, 85 years, female, T1bN0M0- G2L0V0Pn0; (3) Clear cell renal carcinoma, 79 years, female, T1bN0M0-G1L0V0Pn0.

Primary liver tumor patients for single-cell chromatin accessibility sequencing (**Figure 4**) were classified as: (1) Hepatocellular carcinoma, 62 years, male, T2N1M0-G2L1V1Pn0; (2) Hepatocellular carcinoma, 84 years, male, T1bN1M0-G2L0V0Pn0; (3) Hepatocellular carcinoma, 70 years, male, T3N1M0-G3L0V1Pn0.

Primary liver tumor patients for tagmentation-based whole genome bisulfite sequencing (**Figure 5**) were classified as: (1) Hepatocellular carcinoma, 63 years, female, T2N0M0-G2L0V1Pn0; (2) Hepatocellular carcinoma, 42 years, female, T1bN1M0-G1L0V0Pn0; (3) Hepatocellular carcinoma, 78 years, female, T1bN0M0-G2L0V0Pn0.

Metastasis to the liver for single-cell chromatin accessibility sequencing (**Figure 6**) were classified as: (1) Liver metastasis from colorectal adenocarcinoma, 45 years, female; (2) Liver metastasis from colorectal adenocarcinoma, 49 years, male; (3) Liver metastasis from colorectal adenocarcinoma, 57 years, female.

Patient with primary colorectal adenocarcinoma and metastasis to the liver for single-cell chromatin accessibility sequencing (**Figure 7**) was classified as 48 years, female, T3N2bM1b-G3L1V1Pn1.

### Isolation of immune cells from human tissues

For the isolation of immune cells from human tissues, fresh tissue samples were placed in 5 mL digestion buffer per 1 g of tissue and were cut into small pieces before enzymatic digestion on a GentleMACS (Miltenyi Biotec). Digestion buffer and digestion program were tissue type-specific: Colon tissue was digested in RPMI (Gibco #11875093) supplemented with 0.5 mg/mL Collagenase Type II (Sigma-Aldrich #C6885-1G), 50 ug/mL DNase I (Roche #11284932001), 10 mM HEPES (Pan Biotec #P05-01100), and 7.5% FCS (Gibco #26140079) for 60 min and −20rpm at 37°C on the GentleMACS.

Liver tissue was digested in DMEM (Gibco #10938025) containing 1 mg/mL Collagenase Type IV (Sigma-Aldrich #C5138-1G), 50 ug/mL DNase I (Roche #11284932001), 10 mM HEPES (Pan Biotec #P05-01100), and 10% FCS (Gibco #26140079). For liver digestion, the Miltenyi GentleMACS program “h_tumor_01_01” was used, followed by 60 min at 37°C and 45rpm on the GentleMACS. Lung tissue was digested using 1 mg/mL Collagenase Type IV (Sigma-Aldrich #C5138-1G), 50 ug/mL DNase I (Roche #11284932001) in DMEM (Gibco #10938025) containing 5 mg/mL BSA (Serva #11930) for 40 min at 37°C and −20 rpm on a GentleMACS.

Kidney, bladder, and testes were digested using 1 mg/mL Collagenase Type IV (Sigma-Aldrich #C5138-1G), 50 ug/mL DNase I (Roche #11284932001), in DMEM (Gibco #10938025) supplemented with 10 mM HEPES (Pan Biotec #P05-01100) and 10% FCS (Gibco #26140079), for 40 min at 37°C and −20 rpm on a GentleMACS.

Gastric samples were digested in DMEM (Gibco #10938025) containing 2 mg/mL Collagenase Type IV (Sigma-Aldrich #C5138-1G), 20 ug/mL DNase I (Roche #11284932001), 10 mM HEPES (Pan Biotec #P05-01100), and 5 mg/mL BSA (Serva #11930), for 15 min on a rotating shaker at 37°C.

Pancreas tissue was digested with 2 mg/mL Collagenase Type IV (Sigma-Aldrich #C5138-1G), 50 ug/mL DNase I (Roche #11284932001) in DMEM (Gibco #10938025) containing 10 mM HEPES (Pan Biotec #P05-01100) and 5 mg/mL BSA (Serva #11930) for 40 min at 37°C and −20 rpm on a GentleMACS.

Tissue from oral floor was digested in DMEM (Gibco #10938025) supplemented with 4 mg/mL Collagenase Type IV (Sigma-Aldrich #C5138-1G), 50 ug/mL DNase I (Roche #11284932001), 10 mM HEPES (Pan Biotec #P05-01100) and 2% FCS (Gibco Cat#26140079). Digest was performed using the Miltenyi GentleMACS program “37_C_Multi_H”.

Single cells suspensions from all tissues were first filtered through a metal strainer, then a 100 µm strainer followed by a 40 µm strainer prior to enrichment. The single cell suspension was then enriched for CD45^+^ cells using TIL microbeads (Miltenyi Biotec #130-118-780) according to the manufacturer’s protocol.

### Isolation and pre-enrichment of blood lymphocytes

To isolate T cells from human blood, buffy coats were used. Leukocytes were diluted 1:1 with DPBS (Sigma-Aldrich #D8537), and each 20 ml of the resulting mixture was layered onto 20 ml of Histopaque-1077 (Sigma-Aldrich #10771). Samples were centrifuged at 400xg for 30 minutes at RT, with acceleration and brake set to lowest setting. The PBMC layer was isolated and washed three times.

Cells were pre-enriched with PE-labeled anti-human CD25 (Clone BC96, Biolegend), followed by column-based magnetic separation with anti-PE ultrapure microbeads (Miltenyi Biotec #130-105-639) following manufacturer’s protocol.

### FACS sorting

T cells were isolated and pre-enriched as described in the previous sections. Cells were stained in 1.5 mL Eppendorf tubes in FACS buffer (2% FCS in PBS). Surface staining was performed at 4 °C for 20 minutes in 100 µl staining volume. Antibodies were used, if not indicated otherwise, as recommended by the manufacturer. For sorting, the single cell suspension was stained with the following antibodies: CD45- BUV737 (BD Biosciences #748719, clone HI30) or CD45-PE-Cy7 (Biolegend #304016, clone HI30) or CD45-APC-Cy7 (Biolegend #304014, clone HI30), CD3- BV785 (Biolegend #317330, clone OKT3), CD19-APC (Biolegend #302212, clone HIB19), CD206-APC (Biolegend #321110, clone 15-2), CD4-R718 (BD Biosciences #566352, clone SK3), CD8-BB700 (BD Biosciences #566452, clone RPA-T8), CD25-PE (Biolegend #302606, cloneBC96), CD127-BV711 (Biolegend #351328, clone A019D5), CD45RA-BV510 (Biolegend #304142, clone HI100), CD45RO-PE-Cy7 (Biolegend #304230, clone UCHL1), CCR8-BV421 (BD Biosciences #566379, clone 433H), CD39-BV605 (Biolgend #328236, clone A1), PD1-Viobright515 (Miltenyi #130-120-386, clone REA1165), as well as Zombie NIR live/dead dye (Biolegend #423106) or Zombie RED live/dead dye (Biolegend #423110).

BD CS&T beads were used to validate machine functionality, fluorescence spillover compensation was performed with lymphocytes stained with CD4 in the respective colors.

Flow cytometry data were analyzed using BD FlowJo™ (Version 10.6.2), sorting was performed depending on availability with a BD FACSAriaII™ or BD FACSAriaIII™ cell sorter with 85 µm or 100 µm nozzle and 45 psi of pressure. Post-sort quality controls were performed as applicable. Sort strategies are presented in respective **Extended Data Figures**.

### scATAC-seq library preparation and sequencing

Cells were centrifuged (300 g, 5 min, 4°C) and re-suspended in 100 µL PBS (Sigma-Aldrich #D8537) containing 0.04% BSA (Serva #11930). After another centrifugation step (300 g, 5 min, 4°C), supernatant was removed completely and cells were re-suspended in 45 µL chilled lysis buffer (10 mM TRIS-HCL pH7.4 (Sigma #T2194), 10 mM NaCl (Sigma #59222C), 3 mM MgCl (Sigma #M1028), 0.1% Tween-20 (Biocard #1662404), 0.01% Digitonin (Thermo Fisher #BN2006), 1% BSA (Serva #11930) in nuclease-free water (Sigma ‘W4502-10X50ML)). After exactly 2 min, 50 µL wash buffer (10mM TRIS-HCL ph7.4 (Sigma #T2194), 10 mM NaCl (Sigma #59222C), 3 mM MgCl (Sigma #M1028), 0.1% Tween-20 (Biocard #1662404), 1% BSA (Serva #11930) in nuclease-free water (Sigma ‘W4502- 10X50ML)) were added. Nuclei were centrifuged (300 g, 5 min, 4°C) and washed with 45 µL chilled diluted nuclei buffer (10X Genomics, #2000207). After a second centrifugation step (300 g, 5 min, 4°C), nuclei were re-suspended in 7 µL chilled nuclei buffer (10X Genomics, #2000207), and 1 µL of the nuclei was used for counting. 5 µL were used for the transposition reaction, which was performed according to the manufacturer’s protocol (Single-cell ATAC Gel Beads V1.1 and reagents, 10X Genomics #1000175; GEM Chip H, 10X Genomics #1000161), as was the library prep. Libraries for scATAC-seq were sequenced on a NextSeq 500/550, with a paired-end 34-8-16-34 sequencing strategy on a 75-cycle high-output cartridge (Illumina #20024906). Samples were sequenced to a read depth of about 10.000 median high-quality fragments per cell.

### Bioinformatic analysis - general

Software versions and corresponding references, if available, are indicated upon first mention of the respective software. All analyses that used the programming language R were performed using R v4.0.0.

### Pre-processing of scATAC-seq data

Data pre-processing and analysis were performed as described in ^42^. FASTQ files were generated from BCL files using bcl2fastq, and alignment to the reference genome hg38 was performed using *CellRanger ATAC count* (v2.0.0). *ArchR*^43^ (V1.0.1) was used for pre-processing and analysis. For pre-processing, *fragments.tsv* files output by *CellRanger ATAC* were used for generating the count matrix. Barcodes with less than 1000 unique fragments per cell were discarded, as well as barcodes with TSS Enrichment < 10 and barcodes marked as “non-cell” by *CellRanger ATAC*. Dimensionality reduction was performed using the iterative LSI approach implemented in *ArchR*, followed by Louvain clustering. For cell doublet removal, doublet scores were calculated based on synthetic doublets and barcodes with the highest doublet scores were excluded from analysis.

### Single-cell ATAC-seq cell type annotation, peak calling and motif footprints

Manual, cluster-based cell type annotation was performed using gene scores of prior-knowledge marker genes. Gene scores were calculated with *ArchR*, based on the accessibility of gene-coding regions and the corresponding regulatory elements. For the identification of tissue Treg cells, we further calculated z-scores for a published tissue Treg peak signature using the *addDeviationsMatrix* function of *ArchR*. The tissue Treg signature is described in^35^.

Peak calling was performed on pseudobulk replicates, which were created in a sample-aware fashion using *ArchR*’s *addGroupCoverages* function. Peak calling was performed using *MACS2*^63^ (V2.2.6).

Motif footprinting was performed on pseudobulk replicates (see above). For analysing the accessibility of TF motifs between clusters, the *getFootprints* function implemented in *ArchR* was used.

### Computation of Jaccard indices in scATAC-seq datasets

The analyses described in this section were performed using *ArchR* v1.0.1. We extracted binarized peak matrices using *ArchR*’s *getMatrixFromProject* function (useMatrix = ‘PeakMatrix’, binarize = TRUE). To avoid artefacts due to very low cell numbers, we excluded cell types with fewer than 30 cells. We then downsampled each cell population to the same cell number in order to avoid biases arising from different cell numbers. For each cell population, we subsequently computed a binarized pseudobulk vector with one entry for each peak. The pseudobulk value was 1 if a peak was accessible in at least 10% of the cells and 0 otherwise. To avoid biases arising from differences in sequencing depth, we downsampled the “1” values in the pseudobulk vectors to the same number across two compared cell populations. We then computed a Jaccard index between the two downsampled pseudobulk vectors using the *jaccard* function from the *jaccard* R package (v0.1.0)^64^. To investigate how large Jaccard indices between tumor CCR8+ Treg cells and NAT CCR8+ Treg cells would be under the assumption of cell type equality, we simulated 1,000 data sets of tumor CCR8+ Treg cells and NAT CCR8+ Treg cells. In each iteration, the two simulated populations encompassed the same cell number that was used for computation of the actual Jaccard index and were randomly sampled from the same cell type (tumor CCR8+ Treg cells), ensuring that the two simulated populations contained distinct cells. We then repeated the Jaccard index computation as described above, yielding a distribution of expected Jaccard indexes under the assumption of cell type equality.

### Plotting of chromatin accessibility signatures as heatmaps

The analyses described in this section were performed using *ArchR* v1.0.1. We utilized a core Treg cell signature and a tissue Treg cell signature from our previous work^38^. To visualize accessibility in cell types from healthy donors, we utilized the same scATAC-seq data that had been used to generate the signatures^35, 38^. This data set (and hence the signatures) were generated based on the hg19 reference genome whereas the data of tumor patients corresponded to the hg38 reference genome. Thus, we lifted over all peaks from the healthy-donor data set to hg38 using the *CrossMap.py bed* command from *CrossMap*^65^ v0.6.1 and a corresponding chain file downloaded on 03 Jun 2024 from http://hgdownload.soe.ucsc.edu/goldenPath/hg19/liftOver/hg19ToHg38.over.chain.gz; We retained each peak that could be lifted over to a single interval on a normal chromosome (chromosomes 1-22 and chromosome X). Within the tumor-patient data, we then generated peak matrices for the healthy-donor peak set (lifted over to hg38) using *ArchR*’s *addFeatureMatrix* function. To visualize chromatin accessibility in cell types from healthy donors, we used *Signac*^66^ (v1.2.1) and *Seurat*^67^ (v4.0.1). We restricted the raw binarized peak matrix for cells from the relevant cell types using the *GetAssayData* function (slot = ‘counts’). To visualize chromatin accessibility in cell types from tumor patients, we extracted raw binarized peak matrices for healthy-donor peaks lifted over to hg38 using *ArchR*’s *getMatrixFromProject* function (binarize = TRUE). We then concatenated peak matrices for healthy-donor data and tumor-patient data into a joint matrix and normalized data using a term frequency – inverse document frequency appeoach (*RunTFIDF* function from *Signac*)^68^. We aggregated normalized healthy-donor data into cell-type-level values and tumor-patient data into sample-level values by taking the mean across all relevant cells. We extracted values for peaks from the signatures and clipped normalized accessibility values at the 95^th^ percentile. We used the *ComplexHeatmap* package^69^ (v2.6.2) for heatmap generation.

### Statistics related to accessibility heatmaps with respect to individual features

For each tumor patient sample and each signature peak, we compared the methylation value to the respective methylation values in the two healthy-donor cell types shown in the heatmap. We then computed the proportion of signature peaks where accessibility in this sample was closer to the first healthy-donor cell type and the proportion of signature regions where accessibility in this sample was closer to the second healthy-donor cell type. Statistical analysis was performed using Prism software and a two-tailed paired t test.

### Investigating TE accessibility landscapes

The analyses described in this section were performed using ArchR v1.0.1. We utilized RepeatMasker annotations (locations of repeats in the human genome), which had been downloaded from https://hgdownload.soe.ucsc.edu/goldenPath/hg19/database/rmsk.txt.gz on 14 Sep 2023. We restricted healthy-donor data to cells from the relevant cell types that originated from donors that had donated the respective tissue. For these cells, we extracted the raw binarized peak matrix using the *GetAssayData* function (slot = ‘counts’). We extracted raw binarized peak matrices for tumor patient data using *ArchR*’s *getMatrixFromProject* function (binarize = TRUE). Afterwards, we concatenated healthy-donor data and tumor-patient data into a joint peak matrix and normalized this matrix using *Signac*’s *RunTFIDF* function. For the actual analysis, we restricted the data set to peaks from the “Skin Treg hyperaccessibility” signature^38^ that overlapped with relevant TE insertion sites. For each of these peaks and each sample, we computed the mean normalized accessibility across all relevant cells.

### Whole-genome bisulfite sequencing

DNA was isolated using a DNA/RNA micro kit (Qiagen #80284) and DNA quality and concentration was assessed using Qubit^TM^ (dsDNA HS Kit #Q32851) and Tapestation 4200 with Genomic DNA Screentape (Agilent #5067-5365). Tagmentation-based whole genome bisulfite sequencing (TWGBS) was essentially performed as described in literature^36, 37^. Two differently barcoded libraries were generated per sample. Corresponding libraries were pooled in equimolar amounts, and each pool was sequenced on a single lane using Illumina Novaseq 6000 with paired-end sequencing and 150 cycles (PE150).

### Pre-processing of whole-genome bisulfite sequencing data

Alignment and methylation calling was performed based on a described workflow^36^ that employs a customized version of *MethylCtools*^70^. The workflow was modified as follows: *Trimmomatic*^71^ (v0.30) PE was used to trim adaptor sequences with the options “-threads 12 -phred33 ILLUMINACLIP:xxx:2:30:10:8:true SLIDINGWINDOW:4:15 MINLEN:36”. Here, “xxx” refers to a Fasta file containing the adaptor sequences. Alignment to the hs37d5 reference genome (based on GRCh37) was performed using *BWA-MEM*^72^ (v0.7.8) with the “-T” parameter set to 0. Duplicates were marked using *sambamba*^73^ (v0.6.5) with the parameters “-t 1 -l 0 --hash-table-size=2000000 --overflow-list-size=1000000 --io-buffer-size=64” and not removed from the data. For the analysis of CpGs, we combined data on the two opposite cytosines of the same CpG by summing up their read counts. To compute position-wise coverage, we added the counts of methylated and unmethylated reads at each profiled position. We then stored the counts of methylated and total reads at each position in a bsseq object (*bsseq*^74^ package; v1.26.0). We confirmed successful bisulfite conversion in all sequencing libraries by observing low methylation signal at CH sites (**Extended Data Figure 3 and 5**). Here, we restricted our analysis to the first 3,000,000 CH sites on chromosome 1. We computed smoothed methylation values using the *BSmooth* function from the *bsseq* package.

### Analysis methylation across all chromosomes

To visualize methylation across the whole genome, we divided the hs37d5 reference genome into bins of 1,000,000 bases. For each bin and each sample, we then computed the mean raw methylation across all CpGs in that bin using *bsseq*’s *getMeth* function (regions = bins, type = ‘raw’, what = ‘perRegion’). We averaged these bin-level values across all samples of a cell type (*rowMeans* function with “na.rm = TRUE”) and computed the deviation of this value from the average value of blood CD45RA^+^ Tconv cells. We removed bins on the mitochondrial chromosome. To plot genome-wide methylation together with chromosome band information, we utilized the *circos.initializeWithIdeogram* (species = ‘hg19’, plotType = c(’ideogram’, ‘labels’), chromosome.index = relevant_chromosomes) function from *circlize*^75^ (v0.4.13). We then plotted the methylation values using *circlize*’s *circos.track* and *circos.yaxis* functions. To calculate global methylation values for autosomes and chromosome X, we extracted raw sample-level methylation values for every CpG using *bsseq*’s *getMeth* function (type = ‘raw’, what = ‘perBase’). We then computed sample-level mean methylation values across CpGs in the respective chromosome categories (all autosomes or chromosome X) and averaged the resulting values across all samples of a cell type.

### Plotting of DNA methylation signatures as heatmaps

We utilized signatures from our previous work^38^. To visualize methylation in cell types from healthy donors, we utilized the same bisulfite sequencing data that had been used to generate the signatures^38^. For each sample therein and each signature region, we extracted averaged raw methylation values across all recpective CpG sites using *bsseq*’s *getMeth* function (regions = signature_regions, what = ‘perRegion’, type = ‘raw’). We then aggregated these into cell-type-level values by computing the mean across all samples of each cell type. To visualize methylation in cell types from tumor patients, we computed sample-level raw meythylation values for each signature region by averaging methylation across all respective CpG sites (*bsseq*’s *getMeth* function with the parameters “regions = signature_regions, what = ‘perRegion’, type = ‘raw’”). We used the *ComplexHeatmap* package for heatmaps.

### Statistics related to methylation heatmaps with respect to individual features

We considered all features for which all samples from tumor patients had a non-missing methylation value. For each tumor patient sample and each signature region, we compared the methylation value to the respective methylation values in the two healthy-donor cell types shown in the heatmap. We then computed the proportion of signature regions where methylation in this sample was closer to the first healthy-donor cell type and the proportion of signature regions where methylation in this sample was closer to the second healthy-donor cell type. Statistical analysis was performed using Prism and a two-tailed paired t test.

### Investigating TE methylation landscapes

We utilized RepeatMasker annotations (locations of repeats in the human genome), which had been downloaded from https://hgdownload.soe.ucsc.edu/goldenPath/hg19/database/rmsk.txt.gz on 14 Sep 2023. We restricted our analysis to insertion sites of the major TE classes (“DNA”, “LINE”, “LTR” and “SINE”) that overlapped with skin Treg hypomethylation regions from the tissue Treg cell methylation signature^38^. For samples from healthy donors and tumor patients, we extracted sample-level average raw methylation values across all relevant CpG sites using *bsseq*’s *getMeth* function (regions = relevant_insertion_sites, type = ‘raw’, what = ‘perRegion’). To ensure comparability between each sample in the subsequent plot, we excluded insertion sites for which at least one sample displayed a missing value.

### Generation of track plots with methylation data (with or without chromatin accessibility and ChIP-seq data)

To visualize methylation data in selected genomic regions, we utilized track plots. We generated methylation tracks by extracting smoothed methylation values for each CpG and sample using *bsseq*’s *getMeth* function (regions = regions_of_interest, type = ‘smooth’, what = ‘perBase’). The track plot consists of lines connecting CpG-wise values for each cell type or sample. To visualize chromatin accessibility for healthy donors, we utilized Bam files of scATAC-seq data^35^. If necessary, we merged Bam files corresponding to subsets of the same cell type using sambamba merge. We then generated BigWig files containing scATAC-seq signal aggregated into 50-base-pair genomic bins. For this, we employed RPKM normalization using bamCoverage (-of bigwig --normalizeUsing RPKM) from the deeptools^76^ suite (v3.5.1). To visualize chromatin accessibility for tumor patients, we generated BigWig files using *ArchR*’s *getGroupBW* function. Afterwards, we lifted over these BigWig files to hg19 coordinates using the *CrossMap.py bigwig* command with a corresponding chain file downloaded on 03 Dec 2021 from http://hgdownload.soe.ucsc.edu/goldenPath/hg38/liftOver/hg38ToHg19.over.chain.gz; We utilized the import function from the rtracklayer^77^ package (v1.50.0) to load BigWig files into memory. In order to harmonize chromatin accessibility tracks from healthy donors andtumor patients, we linearly scaled the signal from each BigWig file to a range from 0 to 1. We also utilized Chromatin immunoprecipitation followed by sequencing (ChIP-seq) data for BATF in human GM12878 cells^51^, which we obtained from the Gene Expression Omnibus (accession number: GSM803538; downloaded on 09 May 2025). We used the corresponding BigWig file from replicate 2, as in a previous study^35^. To visualize genes in the corresponding regions, we utilized locations of gene bodies and exons in the hs37d5 reference genome that were computed based on the Gencode annotation^78^, version 19. We inferred transcription start sites by the start and end position of gene body intervals together with genomic strand information. For genes on the plus strand, we identified the start of the gene body as the transcription start site. For genes on the minus strand, we identified the end of the gene body as the transcription start site. We aligned all data panels using the *plot_grid* function (ncol = 1, axis = ‘lr’, align = ‘v’) from *cowplot* (Claus O. Wilke (2020). cowplot: Streamlined Plot Theme and Plot Annotations for ‘ggplot2’. R package version 1.1.1. https://CRAN.R-project.org/package=cowplot) (v1.1.1).

### *In-vitro* induction of the tissue Treg program from CD45RA^+^ Treg cells

For *in-vitro* induction of the tissue Treg program, CD45RA^+^ T cells were isolated and pre-enriched as described in the previous sections from buffy coats. The cells were prepared for FACS sorting as described in the earlier sections and stained, if not indicated otherwise, as recommended by the manufacturer. For sorting, the single cell suspension was stained with the following antibodies: CD4-R718 (BD Biosciences #566352, clone SK3), CD19-APC (Biolegend #302212, clone HIB19), CD206-APC (Biolegend #321110, clone 15-2), CD25-PE (Biolegend #302606, cloneBC96), CD127-BV711 (Biolegend #351328, clone A019D5), CD45RA-BV510 (Biolegend #304142, clone HI100), as well as Zombie NIR live/dead dye (Biolegend #423106).

Sorting was performed with a BD FACSAriaII™ cell sorter with 85 µm or 100 µm nozzle and 45 psi of pressure. Post-sort quality controls were performed as applicable. CD45RA^+^ Treg cells were sorted into 15 mL falcons containing 4 mL exMACS^TM^ Medium (Miltenyi Biotec #130-097-196) + 20% FCS (Gibco #26140079).

After sorting cells were centrifuged at 500xg for 5 min at 4°C and seeded in a concentration of 100.000 cells/well in Corning® 96 Well TC-Treated Microplates (Corning #CLS3799-50EA) in TexMACS^TM^ medium supplemented with 1% Penicillin/Streptomycin. The medium was further supplemented for two different conditions:

1. (“Control Treg”): 1x T Cell TransACT^TM^ (Miltenyi Biotec #130-128-758) and 500U IL-2 (Proleukin S, Clinigen PZN 02238131)
2. (“Tissue Treg”): 1x T Cell TransACT^TM^ (Miltenyi Biotec #130-128-758), 500U IL-2 (Proleukin S, Clinigen PZN 02238131), 1x IL-12 (Miltenyi Biotec #130-096-704), 1x IL-21 (Miltenyi Biotec #130-094-563), 1x IL-23 (Miltenyi Biotec #130-095-757), 1x TGFß1 (Miltenyi Biotec #130-095-067)

Cells were fed with another 100 µL of medium accordingly on day 2, and on day 5, half of the medium was replaced. On day 7, 24 h before harvest, the whole medium was replaced with 200 µL of fresh medium. On day 8, the supernatant and cells were harvested. The former was used for the tumor spheroid assay and proteomic analysis, the latter was prepared for FACS analysis. The single cell suspension was stained as described earlier, afterwards the cells were fixed using the eBioscience™ Foxp3 / Transcription Factor Fixation/Permeabilization Concentrate and Diluent Kit (eBiosciences #00-5521-00) according to manufacturer’s instructions. For intracellular staining the following antibodies were used: FoxP3-AF488 (Biolegend #320112), BATF (Cell Signaling Technology #8638S) and Rabbit IgG-AF647 (Biolegend #406414).

### Tumor spheroid assay and image analysis

The human colon cancer cell line HCT116 was used to generate tumor spheroids. 24 hours before harvest of the Treg cell culture, HCT116 were seeded at a concentration of 4000 cells/well in Corning® Costar® Ultra-Low Attachment Multiple Well Plate (Merck #CLS7007-24EA) in 200 µL TexMACS^TM^ medium supplemented with 1% Penicillin/Streptomycin. After confirming the spheroid formation, the spheroids were stimulated by replacing half of the medium with either supernatant from condition 1 (“control Treg”) or condition 2 (“tissue Treg”) of the Treg culture in different dilutions (1:8, 1:16, 1:32, or 1:64 in fresh medium) or fresh medium only.

The spheroids were cultured for 5 days, the plates were placed in the IncuCyte S3 Live Cell Analysis Instrument (Satorius) and scans for each well were scheduled for every 60 mins. The brightfield object total area was calculated with the IncuCyte Spheroid Analysis Software Module (Sartorius) and further analysis were generated with GraphPad Prism (GraphPad Software).

### Quantitative proteomics of tissue Treg supernatant and sorted Treg cells

Proteins derived from cell lysates were processed by single-pot solid-phase-enhanced sample preparation (SP3) as detailed before with minor modifications^79,80^. To this end, cellular lysates were incubated with 20 mM DTT for 30 min at 45 °C to reduce proteins. Afterwards, proteins were alkylated for 30 min at room temperature using iodoacetamide (IAA). Excess IAA was quenched by the addition of DTT. Subsequently, 2 µL of carboxylate-modified paramagnetic beads (Sera-Mag SpeedBeads, GE Healthcare, 0.5 μg solids/μL in water as described in ^79^ were added to the samples. After adding acetonitrile to a final concentration of 70% (v/v), samples were allowed to settle at room temperature for 20 min. Samples were mixed after 10 min. Subsequently, beads were immobilized by incubation on a magnetic rack for 2 min and washed twice with 70% (v/v) ethanol in water and once with acetonitrile. Beads were resuspended in 50 mM NH_4_HCO_3_ supplemented with trypsin (Mass Spectrometry Grade, Promega) at an enzyme-to-protein ratio of 1:25 (w/w) and incubated overnight at 37 °C. After overnight digestion, acetonitrile was added to the samples to reach a final concentration of 95% (v/v). Subsequently, samples were incubated for 20 min at room temperature (and mixed after 10 min). To increase the yield, supernatants derived from this initial peptide-binding step were additionally subjected to the SP3 peptide purification procedure as described before in ^79^. Each sample was washed with acetonitrile. To recover bound peptides, paramagnetic beads from the original sample and corresponding supernatants were pooled in 2% (v/v) dimethyl sulfoxide (DMSO) in water and sonicated for 1 min. After 2 min of centrifugation at 16,200xg and 4 °C, supernatants containing tryptic peptides were transferred into a glass vial for MS analysis and acidified with 0.1% (v/v) formic acid.

### Filter-aided sample preparation (FASP)

Cell culture supernatants were processed by filter-aided sample preparation (FASP). To this end, the protein concentration was determined using the Pierce 660 nm protein assay (Thermo Fisher Scientific) according to the manufactureŕs protocol. 20 µg of total protein were subjected to tryptic digestion using a modified Filter Aided Sample Preparation (FASP) as detailed before in ^81, 82^. In brief, samples (corresponding to approx. 20 µg total protein amount) transferred onto spin filter columns (Nanosep centrifugal devices with Omega membrane, 30 kDa MWCO; Pall, Port Washington, NY). Afterwards, detergents were removed washing the samples (membrane) three times with a buffer containing 8 M urea and 0.1 M TRIS Base. After reduction and alkylation by DTT and iodoacetamide (IAA), excess IAA was quenched with DTT and the membrane washed three times with 50 mM NH_4_HCO_3_. Afterwards, proteins were digested overnight at 37° C with trypsin (Trypsin Gold, Promega, Madison, WI) using an enzyme-to-protein ratio of 1:50 (w/w). After digestion, peptides were recovered by centrifugation and one additional wash with 50 mM NH_4_HCO_3_. Combined flow-throughs were acidified with trifluoroacetic acid (TFA) to a final concentration of 1% (v/v) TFA and lyophilized. Purified peptides were reconstituted in 0.1% (v/v) formic acid (FA) for LC-MS analysis.

### Liquid chromatography-mass spectrometry (LC-MS) analysis

Peptides derived from the cellular digests were separated on a nanoElute LC system (Bruker Corporation, Billerica, MA, USA) at 400 nL/min using a reversed phase C18 column (Aurora UHPLC emitter column, 25 cm x 75 µm 1.6 µm, IonOpticks) which was heated to 50°C. Peptides were loaded onto the column in direct injection mode at 600 bar. Mobile phase A was 0.1% FA (v/v) in water and mobile phase B 0.1% FA (v/v) in ACN. Peptides were separated running a linear gradient from 2% to 37% mobile phase B over 39 min. Afterwards, column was rinsed for 5 min at 95% B. Eluting peptides were analyzed in positive mode ESI-MS using parallel accumulation serial fragmentation enhanced data-independent acquisition mode (diaPASEF) on a timsTOF SCP mass spectrometer (Bruker Corporation) ^83^. The dual TIMS was operated at a fixed duty cycle close to 100% using equal accumulation and ramp times of 100 ms each spanning a mobility range from 1/K_0_ = 0.7 Vs cm^−2^ to 1.3 Vs cm^−2^. We defined 27 × 25 Th isolation windows from *m/z* 325 to 990 resulting in ten diaPASEF scans per acquisition cycle. The collision energy was ramped linearly as a function of the mobility from 59 eV at 1/K_0_ = 1.6 Vs cm^−2^ to 20 eV at 1/K_0_ = 0.6 Vs cm^−2^.

Reconstituted peptides (FASP-digested cell culture supernatants) were analyzed by LC-MS using an Evosep One chromatography system (Evosep, Odense, Denmark) coupled online to a timsTOF HT mass spectrometer (Bruker Corporation, Billerica, MA, USA). For MS-analysis, samples (corresponding to 250 ng of peptides) were loaded onto EvoTips according to the manufactureŕs protocol. Subsequently, peptides were separated using an 8 cm x 150 µm, 1.5 µm reversed phase C18 column (EV1109, Evosep performance column) and the “60 samples per day (60SPD)” method provided by the manufacturer. Column was heated to 40° C. Eluting peptides were analyzed in positive mode ESI-MS using in diaPASEF mode as detailed in ^83^. The dual TIMS (trapped ion mobility spectrometer) was operated at a fixed duty cycle close to 100% using equal accumulation and ramp times of 100 ms each spanning a mobility range from 1/K_0_ = 0.6 Vs cm^−2^ to 1.6 Vs cm^−2^. The diaPASEF scheme was generated using pydiAID covering a precursor mass range from *m/z* 270 to 1,400 ^84^. We defined 24 windows of variable width resulting in 8 diaPASEF scans per acquisition cycle comprising three isolation windows each. The collision energy was ramped linearly as a function of the mobility from 59 eV at 1/K_0_ = 1.3 Vs cm^−2^ to 20 eV at 1/K_0_ = 0.85 Vs cm^−2^. All samples were analyzed in three technical replicates.

### Data analysis and label-free quantification

MS raw data were processed using DIA-NN (version 1.9.2)^85^ applying the default parameters for library-free database search. Data were searched using a custom compiled database containing UniProtKB/SwissProt entries of the human reference proteome and a list of common contaminants (version release March 2024, 20,436 entries). For peptide identification and in-silico library generation, trypsin was set as protease allowing one missed cleavage. Carbamidomethylation was set as fixed modification, and the maximum number of variable modifications was set to zero. The peptide length ranged between 7–30 amino acids. The precursor *m/z* range was set to 300–1,800, and the product ion *m/z* range to 200–1,800. As quantification strategy we applied the “QuantUMS (high accuracy)” mode with RT-dependent median-based cross-run normalization enabled. We used the build-in algorithm of DIA-NN to automatically optimize MS2 and MS1 mass accuracies and scan window size. Peptide precursor FDRs were controlled below 1%. In the final proteome datasets, proteins had to be identified by at least two peptides.

### Identification of proteins secreted by induced tissue Treg cells

To identify proteins that are more strongly secreted by induced tissue Treg cells than by control Treg cells, we utilized a conservative approach demanding that a protein must both display stronger expression on the RNA level and higher levels in proteomics data. To examine the RNA level, we utilized differential expression results based on bulk RNA-seq data of the same cell types^35^. We extracted all genes that had been significantly (Adj.P < 0.05) overexpressed in one of the two compared cell types. We matched protein IDs and gene symbols using an overview table from the UniProt data base (https://www.uniprot.org; Section “Core Data”, subsection “Proteins (UniProtKB)”, subsubsection “Human”; downloaded on 24 Oct 2024). In the subsequent analysis, we considered all genes with significant differential expression that could be unambiguously matched to a single protein in the proteomics data. We then visualised Log2(fold change) values on the RNA level together with Log2(fold change) values on the protein level.

### Mapping of secreted proteins to a gene regulatory network

To obtain a gene regulatory network, we downloaded the CollecTRI resource^86^ of transcription factor target genes uding R v4.4.0 and the *get_collectri* function from the *decoupleR* package^87^ (v2.10.0). We restricted this list to regulatory interactions with target genes that are up-regulated in induced tissue Treg cells both on the RNA level and on the protein level (quadrant 1). Using *igraph*, we subsequently generated a directed graph of these regulatory interactions in which each transcription factor is linked to its target genes. We employed *igraph*’s *degree* function to determine the 10 transcription factors with the highest degree (*i.e.* with the highest number of connected edges). We then restricted the graph to these top-10 transcription factors and the vertices that were directly connected to them (*igraph*’s *delete.vertices* function).

### Computation of Transcript per Million (TPM) values

We converted previously computed reads per kilobase per million (RPKM) values^35^ to transcripts per million (TPM) values by multiplying each RPKM value by 1,000,000 and then dividing by the sum of RPKM values across all genes in the respective sample.

## DATA AND CODE AVAILABILITY

The accession numbers for human scATAC-seq and WGBS data reported in this paper are: European Genome Phenome Archive (EGA) XXXXXXXX. The accession numbers for proteomics data reported in this paper are: XXXXXXXX.

## EXTENDED DATA TABLES, FIGURES AND LEGENDS

**Extended Data Table 1.** Signatures used in this paper.

**Extended Data Table 2.** Factors identified in secretome/gene expression correlation

**Extended Data Figure 1.**
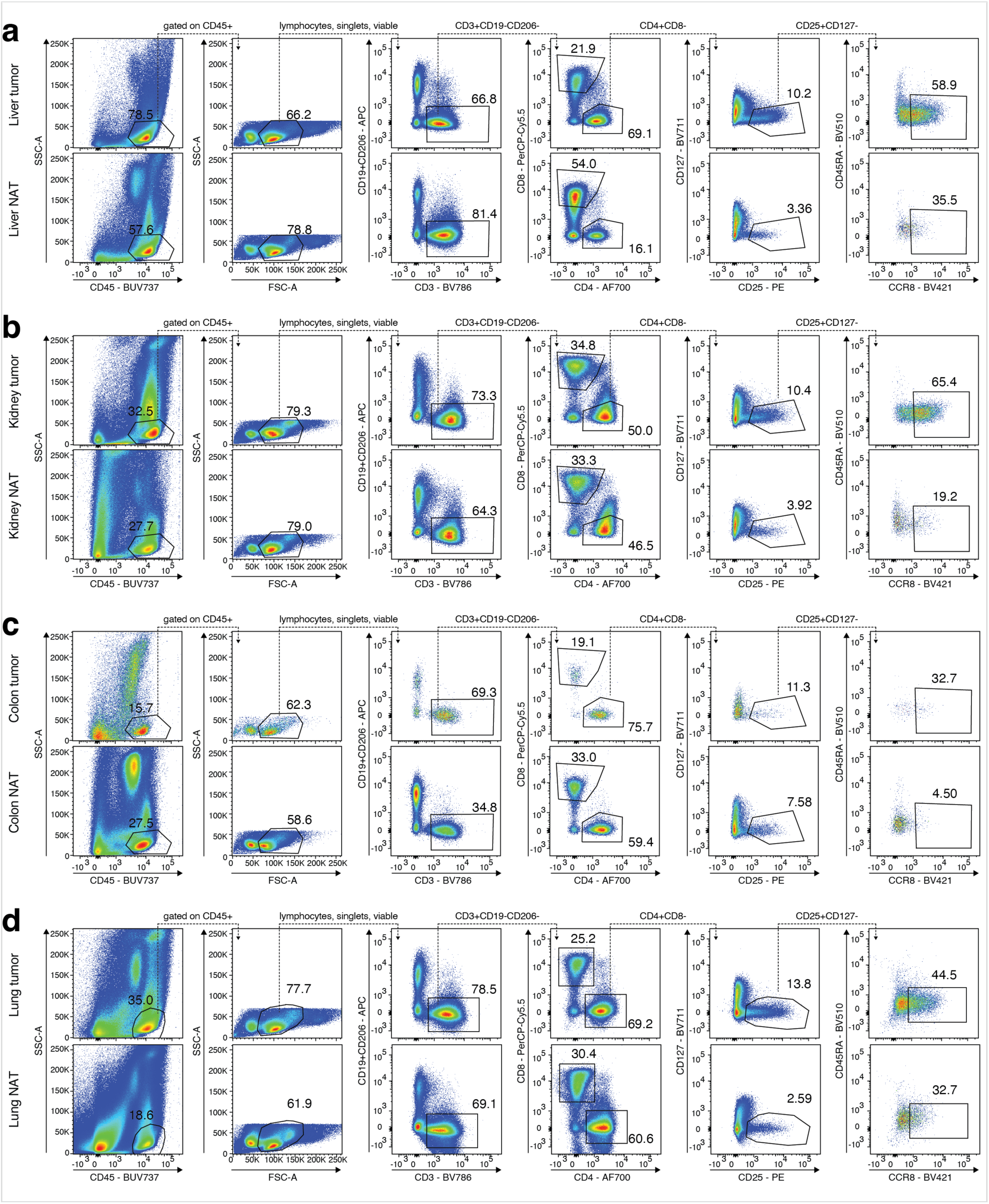
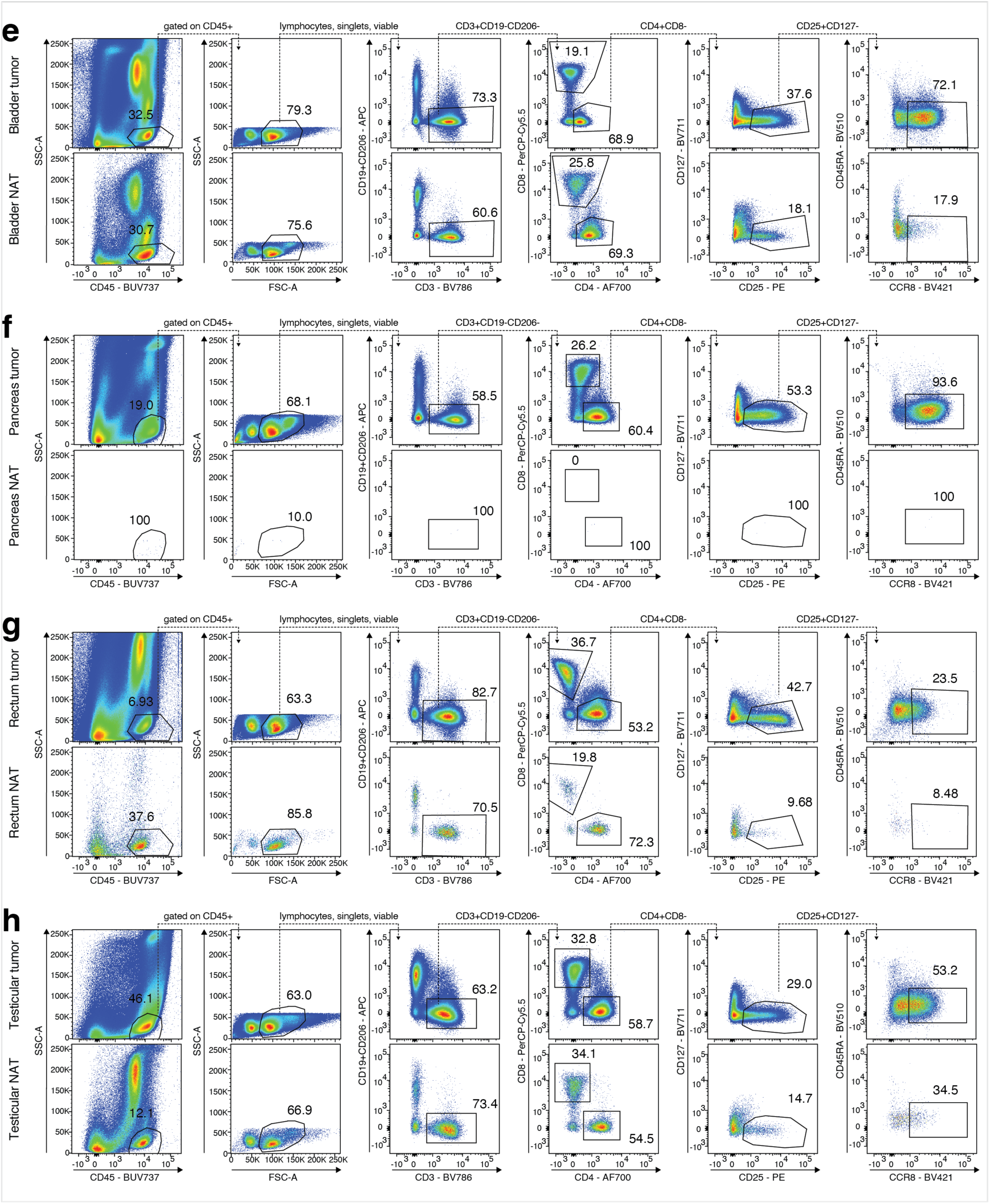
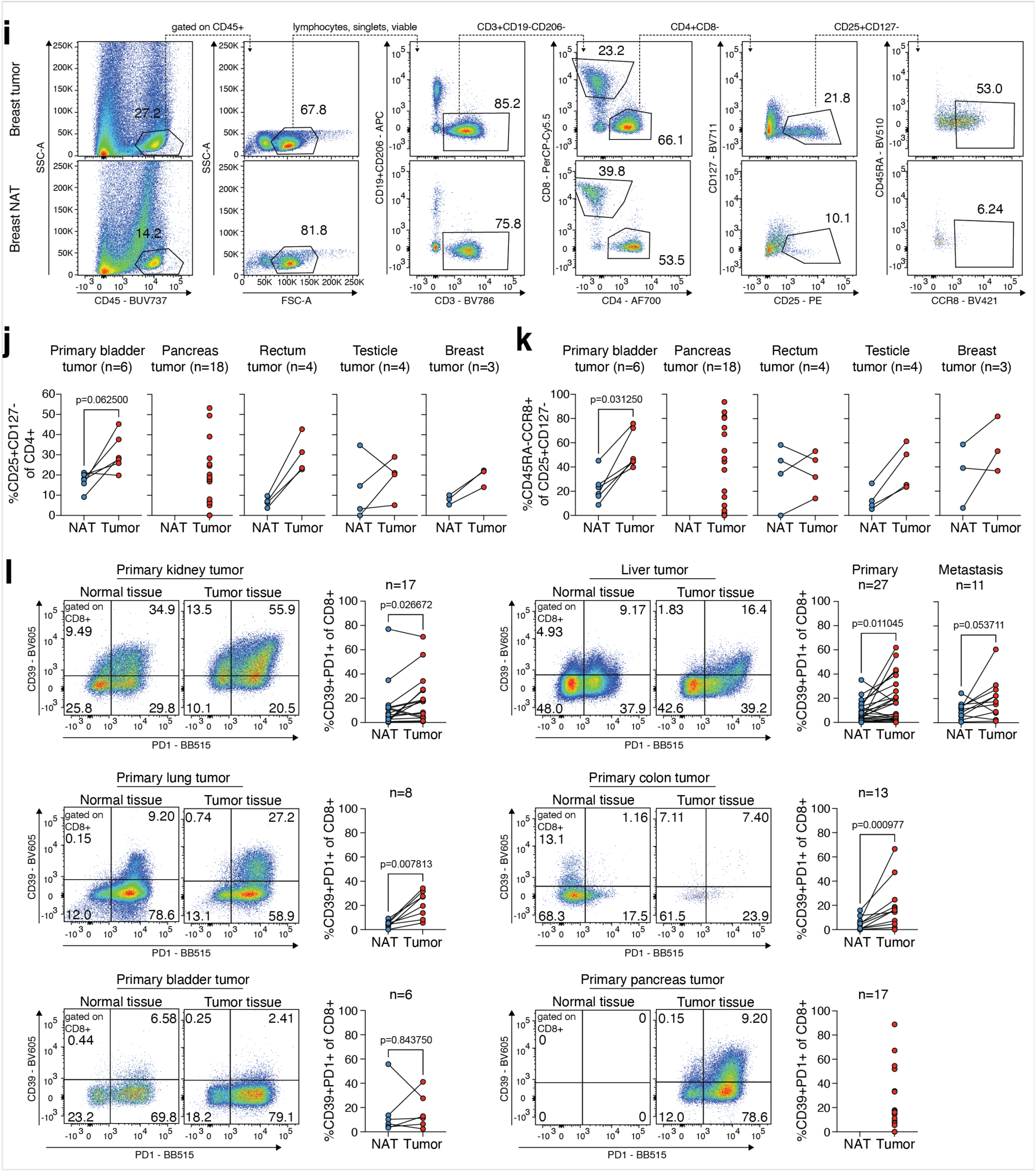
Gating strategies to identify T cell subpopulations from tumor and cognate NAT. **(a-i)** Representative flow cytometry dot plots illustrating the gating of Treg cells (CD127^-^ CD25^high^ CD4^+^ T cells) and CCR8^+^ Treg cells (CD45RA^-^CCR8^+^ from CD127^-^CD25^high^ Treg cells) in tumor and NAT tissue of primary liver tumor or liver metastasis patients (a), primary kidney tumor patients (b), primary colon tumor patients (c), primary lung tumor patients (d), primary bladder tumor patients (e), primary pancreas tumor patients (f), primary rectum tumor patients (g), primary testicular cancer patients (h), and primary breast tumor patients (i), related to Figure 1. **(j-k)** Treg cell frequency of CD4^+^ T cells (j) and CCR8^+^ Treg cell frequency of Treg cells (k) for tumor and healthy tissue isolated from human bladder (n=6), human pancreas (n=18), human rectum (n=4), human testicle (n=4) and human breast (n=3) tumor patients, statistical analysis via Wilcoxon matched-pairs signed-rank test if applicable. **(l)** Representative flow cytometry dot plots illustrating the gating of CD8 Texh cells as CD39^+^PD1^+^ subpopulation of CD8^+^ T cells in tumor and NAT tissue of primary kidney tumor patients (n=17), primary liver tumor (n=27) or liver metastasis patients (n=11), primary lung tumor patients (n=8), primary colon tumor patients (n=13), primary bladder tumor patients (n=6), and primary pancreas tumor patients (n=17), statistical analysis via Wilcoxon matched-pairs signed-rank test. For pancreas, only very low quantities of immune cells were detected in NAT. Data representative of several independent experiments with the indicated number of patients.

**Extended Data Figure 2.**
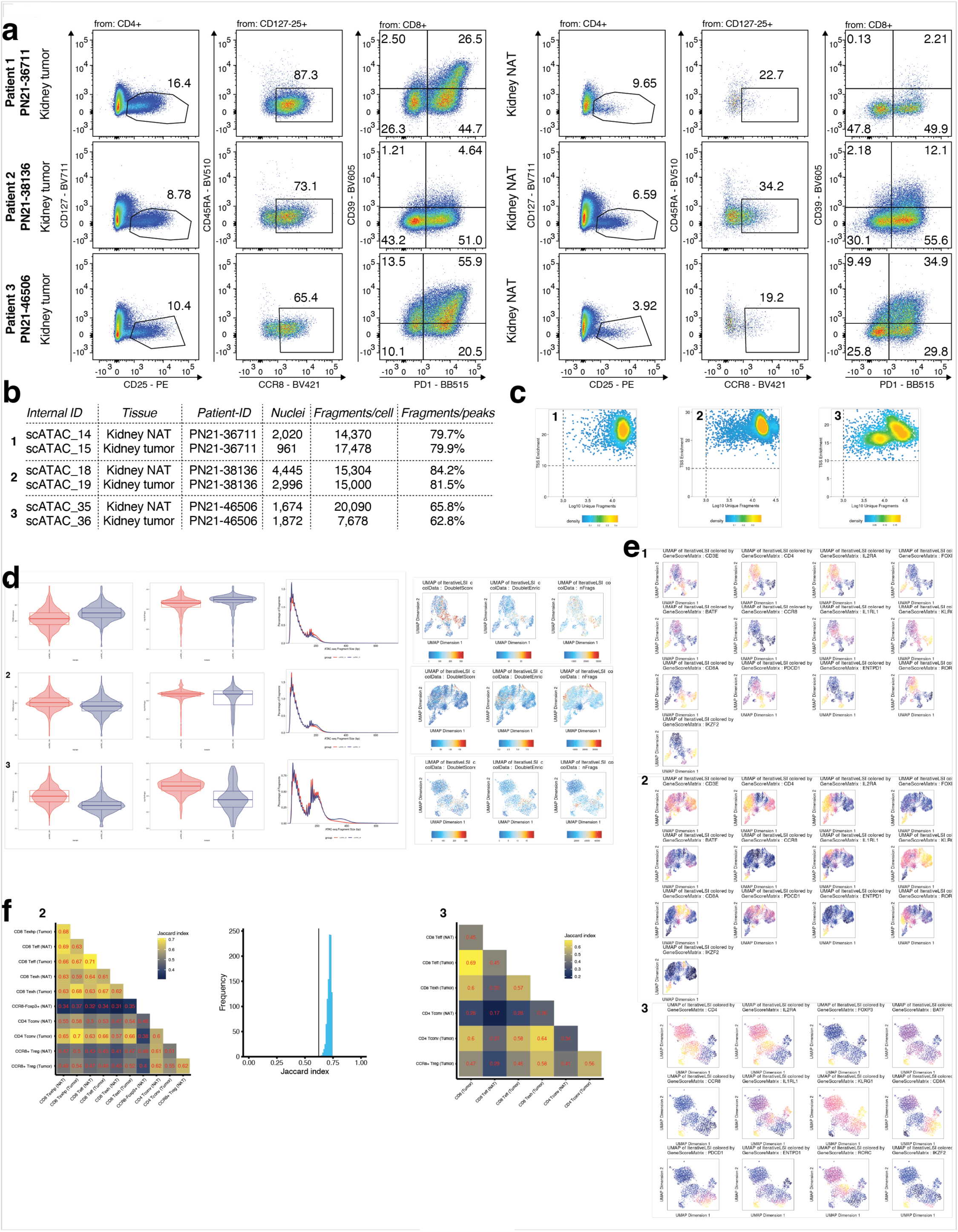
Single-cell ATAC-seq of CD3^+^ T cells from kidney tumor vs NAT. **(a)** Flow cytometry of tumor and NAT of three kidney tumor patients used for scATAC-seq, related to Figure 2. Frequency of CD25^+^CD127^-^ Treg cells of CD4^+^ T cells; CCR8^+^CD45RA^-^ tissue Treg cells of CD25^+^CD127^-^ Treg cells; PD1^+^CD39^+^ Texh cells of CD8^+^ T cells. Left 3 plots, results for tumor; right 3 plots, results for NAT. Precise gating strategy for kidney samples shown in **Extended Data Figure 1b**. **(b)** Results of scATAC-seq *CellRanger ATAC* run for 6 runs with 3 patients. Columns show values for internal ID, tissue type, patient-ID, # of nuclei detected by *CellRanger ATAC*, number of median high-quality fragments per cell, and fraction of high-quality fragments overlapping peaks. **(c)** Dot plot illustrating the TSS enrichment vs Log10 unique fragments for patient 1-3, after data pre-processing using *ArchR*. **(d)** Further quality control after data pre-processing using *ArchR*. Left, TSS Enrichment per cell and log10 number of fragments per cell for NAT (red) vs tumor (blue). Middle, histogram depicting the fragment length distribution in basepairs (bp) for NAT (red) vs tumor (blue). Right, UMAP illustrating the doublet score, double enrichment and number of fragments per cell. **(e)** Imputation of marker gene accessibility scores using *MAGIC* smoothing for *CD3E*, *CD4*, *IL2RA*, *FOXP3*, *BATF*, *CCR8*, *IL1RL1*, *KLRG1*, *CD8A*, *PDCD1*, *ENTPD1*, *RORC*, and *IKZF2*. **(f)** Top, Jaccard indices underlying the mean values presented in Figure 2h. Only cell types with sufficient cell numbers for Jaccard index computation are included. Middle, distribution of expected Jaccard indices assuming equality between Tumor CCR8+ Treg cells and NAT CCR8+ Treg cells. Vertical lines mark the observed Jaccard index between these two populations. Data are only shown for patients for whom a Jaccard index between the two populations could be computed. Data representative of three independent experiments with three individual kidney tumor patients.

**Extended Data Figure 3.**
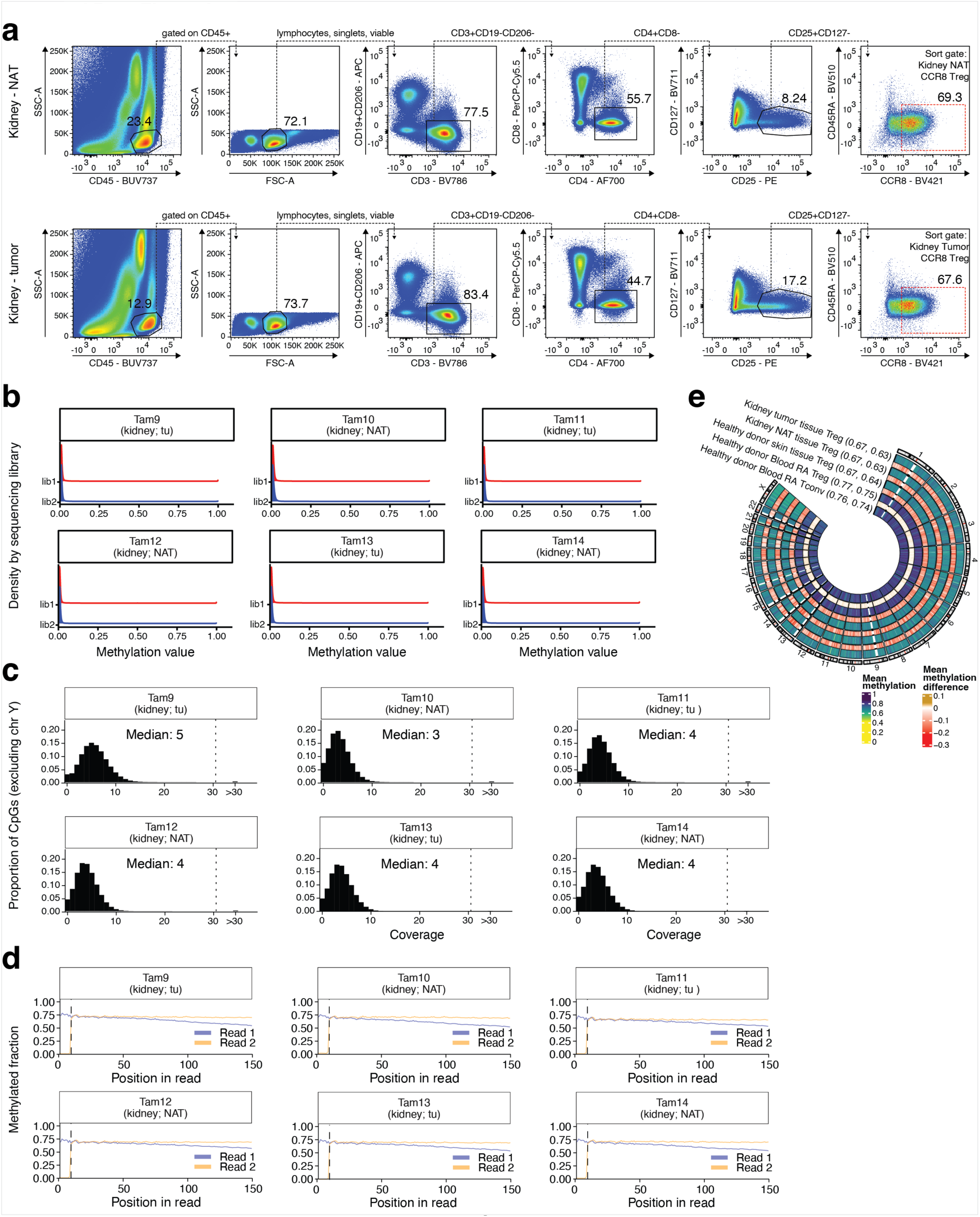
Whole-genome bisulfite sequencing of CCR8^+^ Treg cells from kidney tumor vs NAT. **(a)** Representative flow cytometry dot plots and gating strategy to sort CCR8^+^ Treg cells (CD45RA^-^CCR8^+^ from CD127^-^CD25^+^ Treg cells) in tumor and NAT tissue of primary kidney tumor patients, related to Figure 3. **(b)** Distribution of CH methylation values, stratified by sample and sequencing library, with methylation value on x-axis, for sequencing library 1 (lib1) and sequencing library 2 (lib2). **(c)** Distribution og CpG coverage (excluding the Y chromosome) for whole-genome bisulfite sequencing samples, with coverage on x-axis and median coverage indicated on top. **(d)** M-bias plot showing dependency of methylation on the position in a bisulfite sequencing read, stratified by read type and sample. Vertical dashed lines indicate position 10. **(e)** Methylation level by genomic position and corresponding differences with respect to healthy-donor blood CD45RA^+^ Tconv cells. Numbers in brackets indicate the average methylation level on autosomes and chromosome X, respectively. RA, CD45RA. Data representative of three independent experiments with three individual primary kidney tumor patients.

**Extended Data Figure 4.**
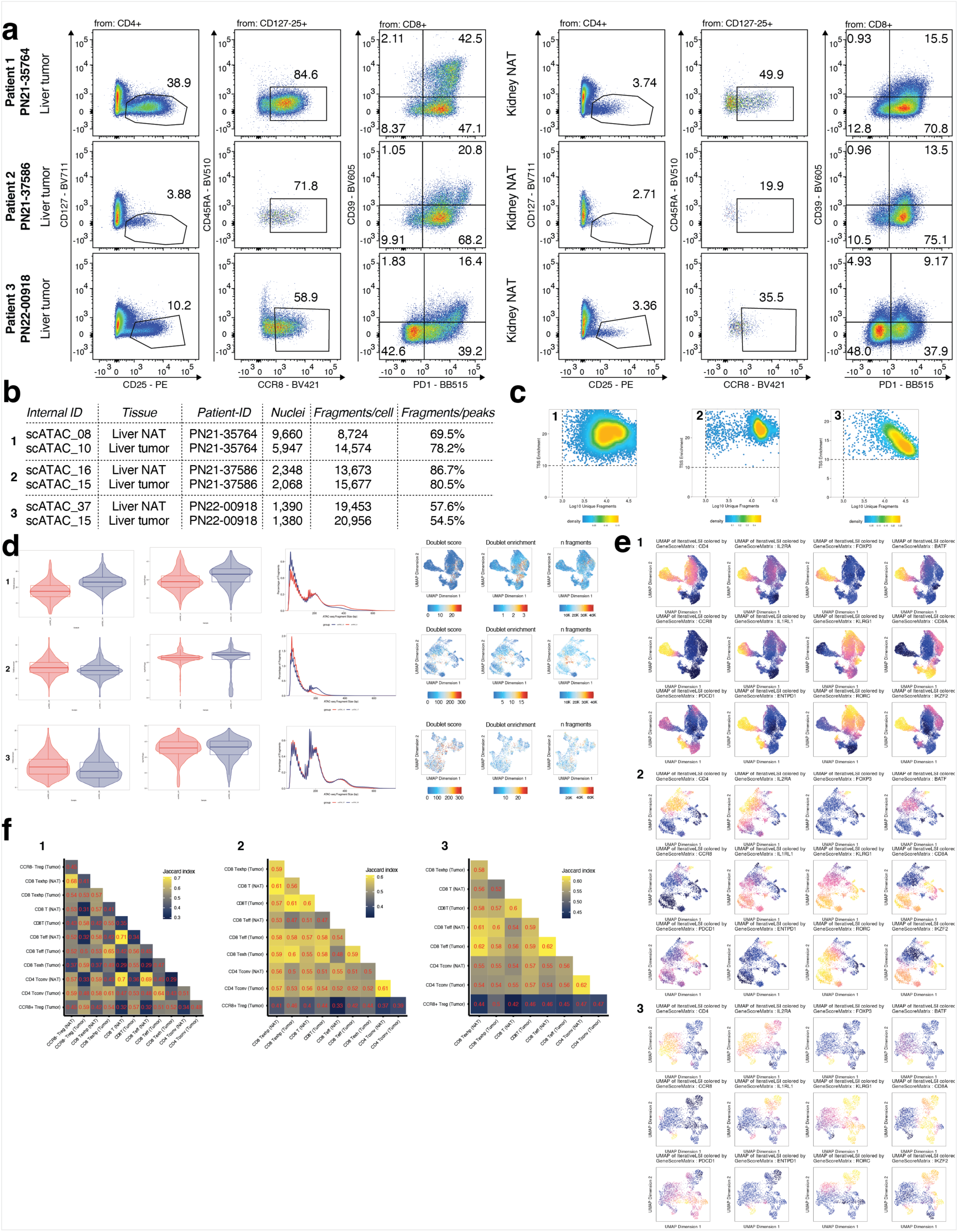
Single-cell ATAC-seq of CD3^+^ T cells from primary liver tumor vs NAT. **(a)** Flow cytometry of tumor and NAT of three primary liver tumor patients used for scATAC-seq, related to Figure 4. Frequency of CD25^+^CD127^-^ Treg cells of CD4^+^ T cells; CCR8^+^CD45RA^-^ tissue Treg cells of CD25^+^CD127^-^ Treg cells; PD1^+^CD39^+^ Texh cells of CD8^+^ T cells. Left 3 plots, results for tumor; right 3 plots, results for NAT. Precise gating strategy for liver samples shown in **Extended Data Figure 1a. (b)** Results of scATAC-seq *CellRanger ATAC* run for 6 runs with 3 patients. Columns show values for internal ID, tissue type, patient-ID, # of nuclei detected by *CellRanger ATAC*, number of median high-quality fragments per cell, and fraction of high-quality fragments overlapping peaks. **(c)** Dot plot illustrating the TSS enrichment vs Log10 unique fragments for patient 1-3, after data pre-processing using *ArchR*. **(d)** Further quality control after data pre-processing using *ArchR*. Left, TSS Enrichment per cell and log10 number of fragments per cell for NAT (red) vs tumor (blue). Middle, histogram depicting fragment length distribution in basepairs (bp) for NAT (red) vs tumor (blue). Right, UMAP illustrating the doublet score, double enrichment and number of fragments per cell. **(e)** Imputation of marker gene accessibility scores using *MAGIC* smoothing for *CD3E*, *CD4*, *IL2RA*, *FOXP3*, *BATF*, *CCR8*, *IL1RL1*, *KLRG1*, *CD8A*, *PDCD1*, *ENTPD1*, *RORC*, and *IKZF2*. (**f**) Jaccard indices between accessible peaks in the analyzed cell types. Only cell types with sufficient cell numbers for Jaccard index computation are included. Data representative of three independent experiments with three individual primary liver tumor patients.

**Extended Data Figure 5.**
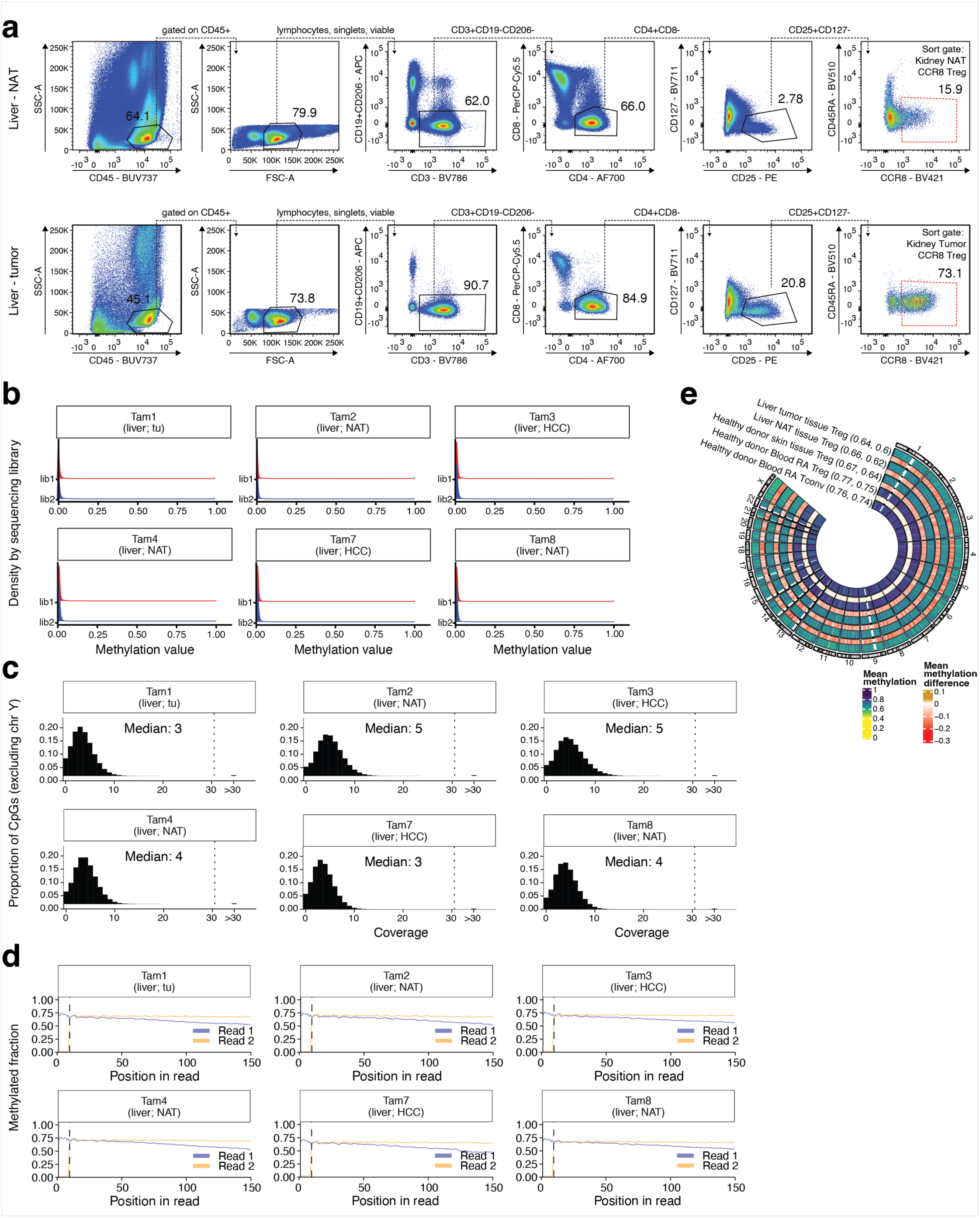
Whole-genome bisulfite sequencing of CCR8^+^ Treg cells from liver tumor vs NAT. **(a)** Representative flow cytometry dot plots and gating strategy to sort CCR8^+^ Treg cells (CD45RA^-^CCR8^+^ from CD127^-^CD25^+^ Treg cells) in tumor and NAT tissue of primary liver tumor patients, related to Figure 5. **(b)** Distribution of CH methylation values stratified by whole-genome bisulfite sequencing library, with methylation value on x-axis, for sequencing library 1 (lib1) and sequencing library 2 (lib2). **(c)** Distribution of CpG coverage values (excluding the Y chromosome) for whole-genome bisulfite sequencing samples, with coverage on x-axis and median coverage indicated on top. **(d)** M-bias plot showing dependency of methylation on the position inside a bisulfite sequencing read, stratified by read type and sample. Vertical dashed lines indicate position 10. **(e)** Methylation level by genomic position and corresponding differences with respect to blood CD45RA^+^ Tconv cells. Numbers in brackets indicate the average methylation level on autosomes and chromosome X, respectively. RA, CD45RA. Data representative of three independent experiments with three individual primary liver tumor patients.

**Extended Data Figure 6.**
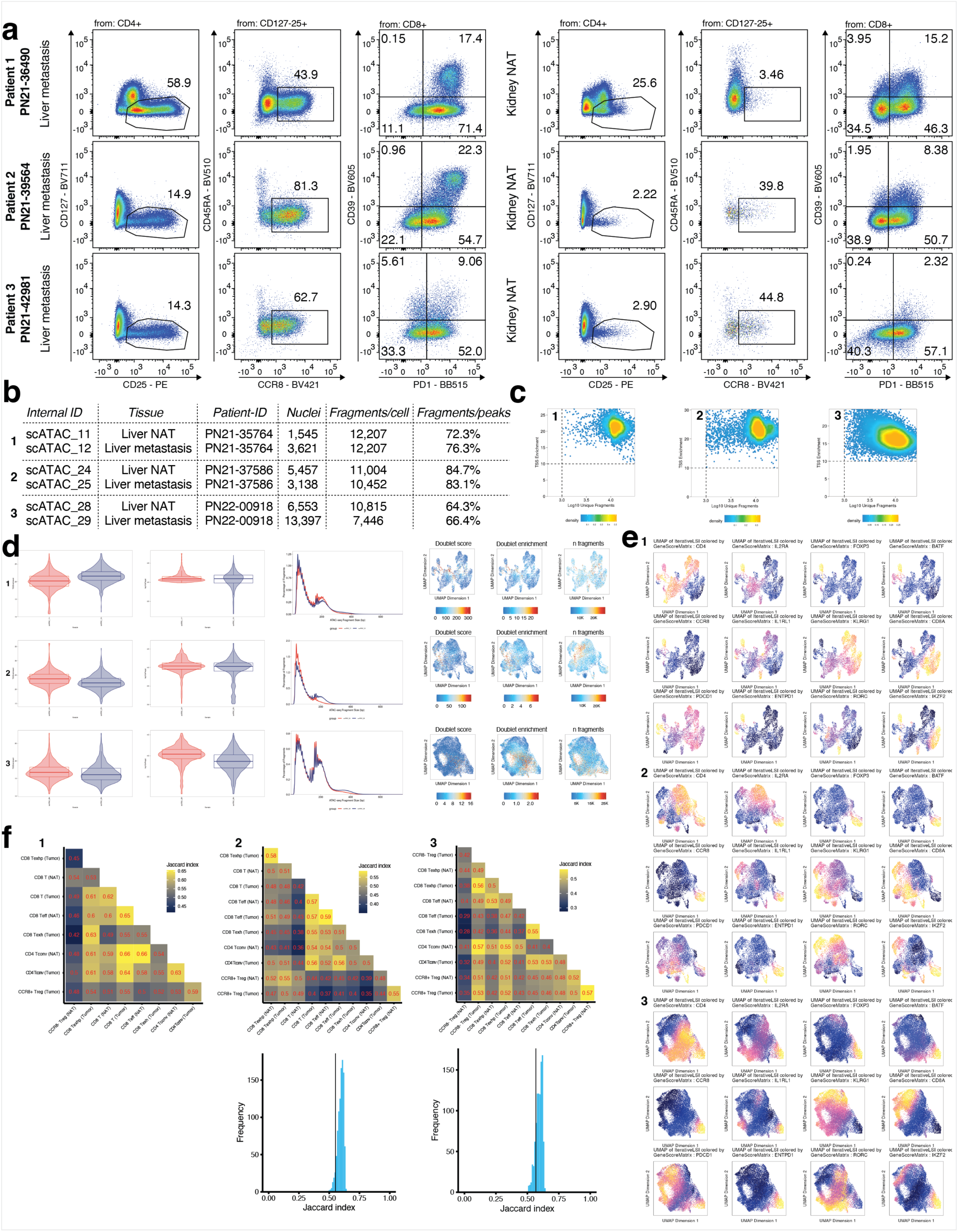
Single-cell ATAC-seq of CD3^+^ T cells from liver metastasis vs NAT. **(a)** Flow cytometry of tumor and NAT of three liver metastasis tumor patients used for scATAC-seq, related to Figure 6. Frequency of CD25^+^CD127^-^ Treg cells of CD4^+^ T cells; CCR8^+^CD45RA^-^ tissue Treg cells of CD25^+^CD127^-^ Treg cells; PD1^+^CD39^+^ Texh cells of CD8^+^ T cells. Left 3 plots, results for tumor; right 3 plots, results for NAT. Precise gating strategy for liver samples shown in **Extended Data Figure 1a**. **(b)** Results of scATAC-seq *CellRanger ATAC* run for 6 runs with 3 patients. Columns show values for internal ID, tissue type, patient-ID, # of nuclei detected by *CellRanger ATAC*, number of median high-quality fragments per cell, and fraction of high-quality fragments overlapping peaks. **(c)** Dot plot illustrating the TSS enrichment vs Log10 unique fragments for patient 1-3, after data pre-processing using *ArchR*. **(d)** Further quality control after data pre-processing using *ArchR*. Left, TSS Enrichment per cell and log10 number of fragments per cell for NAT (red) vs tumor (blue). Middle, histogram depicting fragment length distribution in basepairs (bp) for NAT (red) vs tumor (blue). Right, UMAP illustrating the doublet score, double enrichment and number of fragments per cell. **(e)** Imputation of marker gene accessibility scores using *MAGIC* smoothing for *CD3E*, *CD4*, *IL2RA*, *FOXP3*, *BATF*, *CCR8*, *IL1RL1*, *KLRG1*, *CD8A*, *PDCD1*, *ENTPD1*, *RORC*, and *IKZF2*. **(f)** Top, Jaccard indices underlying the mean values presented in Figure 6h. Only cell types with sufficient cell numbers for Jaccard index computation are included. Bottom, Distribution of expected Jaccard indices assuming equality between Tumor CCR8+ Treg cells and NAT CCR8+ Treg cells. Vertical lines mark the observed Jaccard index between these two populations. Data are only shown for patients for whom a Jaccard index between the two populations could be computed. Data representative of three independent experiments with three individual liver metastasis patients.

**Extended Data Figure 7.**
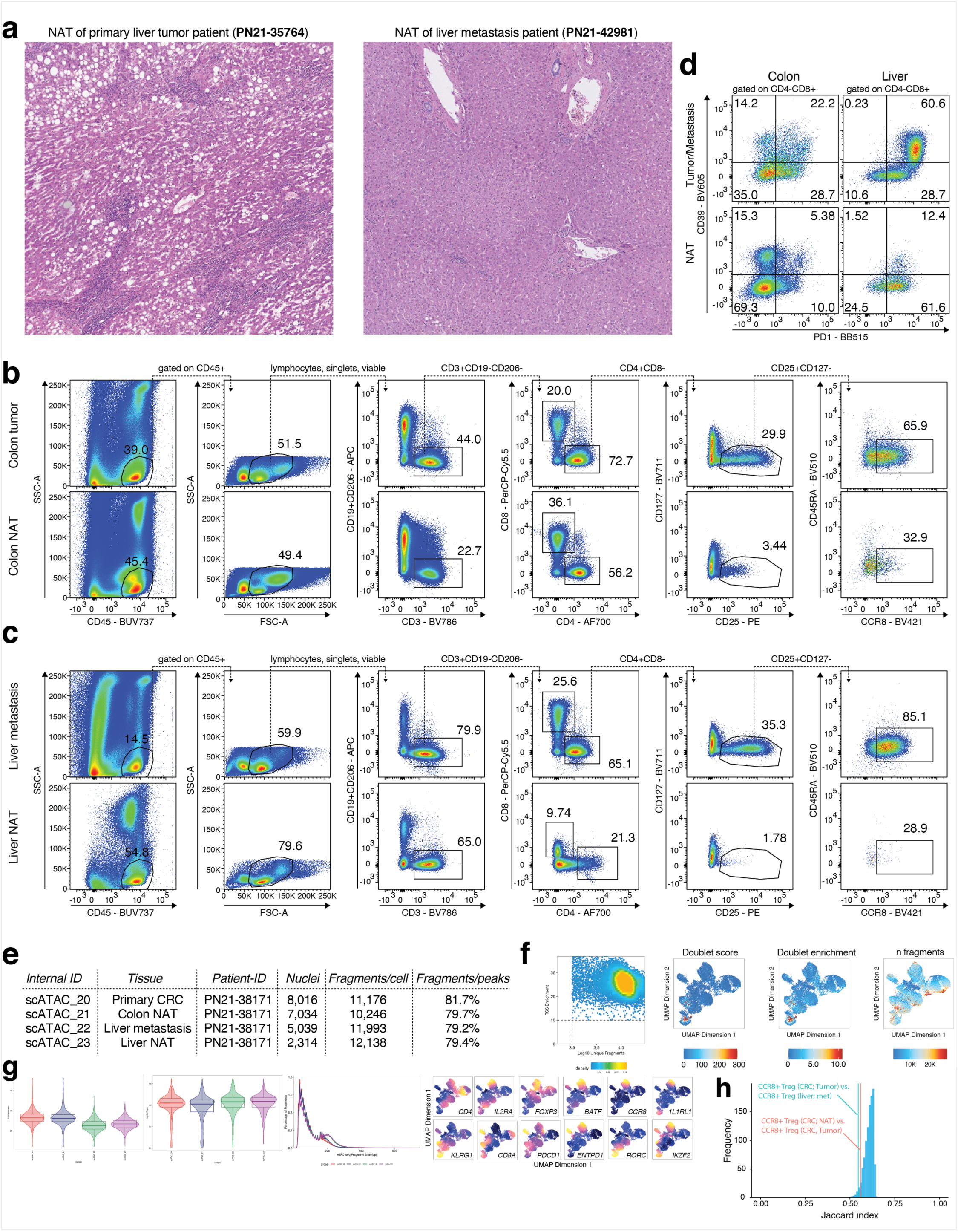
Histology and single-cell ATAC-seq of CD3^+^ T cells from primary colon tumor & NAT and liver metastasis & NAT. **(a)** Histology of NAT from primary liver tumor vs NAT from liver metastasis, related to Figure 6. Left, NAT of patient (primary liver tumor patient 1) shows inflammatory infiltrate (portal and intralobular) as well as steatosis. The lobular structure is lost. Right, NAT of patient (liver metastasis patient 3) shows lobularly structured liver parenchyma with a neglectable degree of inflammatory infiltrate and no steatosis. **(b-c)** Flow cytometry dot plots illustrating the frequency and gating of Treg cells (CD127^-^CD25^high^ CD4^+^ T cells) and CCR8^+^ Treg cells (CD45RA^-^ CCR8^+^ from CD127^-^CD25^high^ Treg cells) in primary colon tumor and cognate colon NAT (b) or in metastasis to the liver from a primary colon tumor and cognate liver NAT as in (c) for patient PN21- 38171, related to Figure 7**. (d)** Frequency of PD1^+^CD39^+^ Texh cells of CD8^+^ T cells in in primary colon tumor and cognate NAT as in (b) as well as metastasis to the liver from a primary colon tumor and cognate liver NAT as in (c) for patient PN21-38171. **(e)** Results of scATAC-seq *CellRanger ATAC* run for 4 runs with patient PN21-38171. Columns show values for internal ID, tissue type, patient-ID, # of nuclei detected by *CellRanger ATAC*, number of median high-quality fragments per cell, and fraction of high-quality fragments overlapping peaks. **(f)** Dot plot illustrating TSS enrichment vs Log10 unique fragments and UMAP illustrating the doublet score, double enrichment and number of fragments per cell. **(g)** TSS Enrichment per cell, log10 number of fragments per cell (left), fragment size distribution (middle), and imputation of marker gene accessibility scores using *MAGIC* smoothing (right). **(h)** Distribution of expected Jaccard indices assuming equality between different Treg cell populations. Vertical lines indicate observed Jaccard indices for two cell type comparisons. Data representative of a single experiment with one tumor and metastasis from one patient.

**Extended Data Figure 8.**
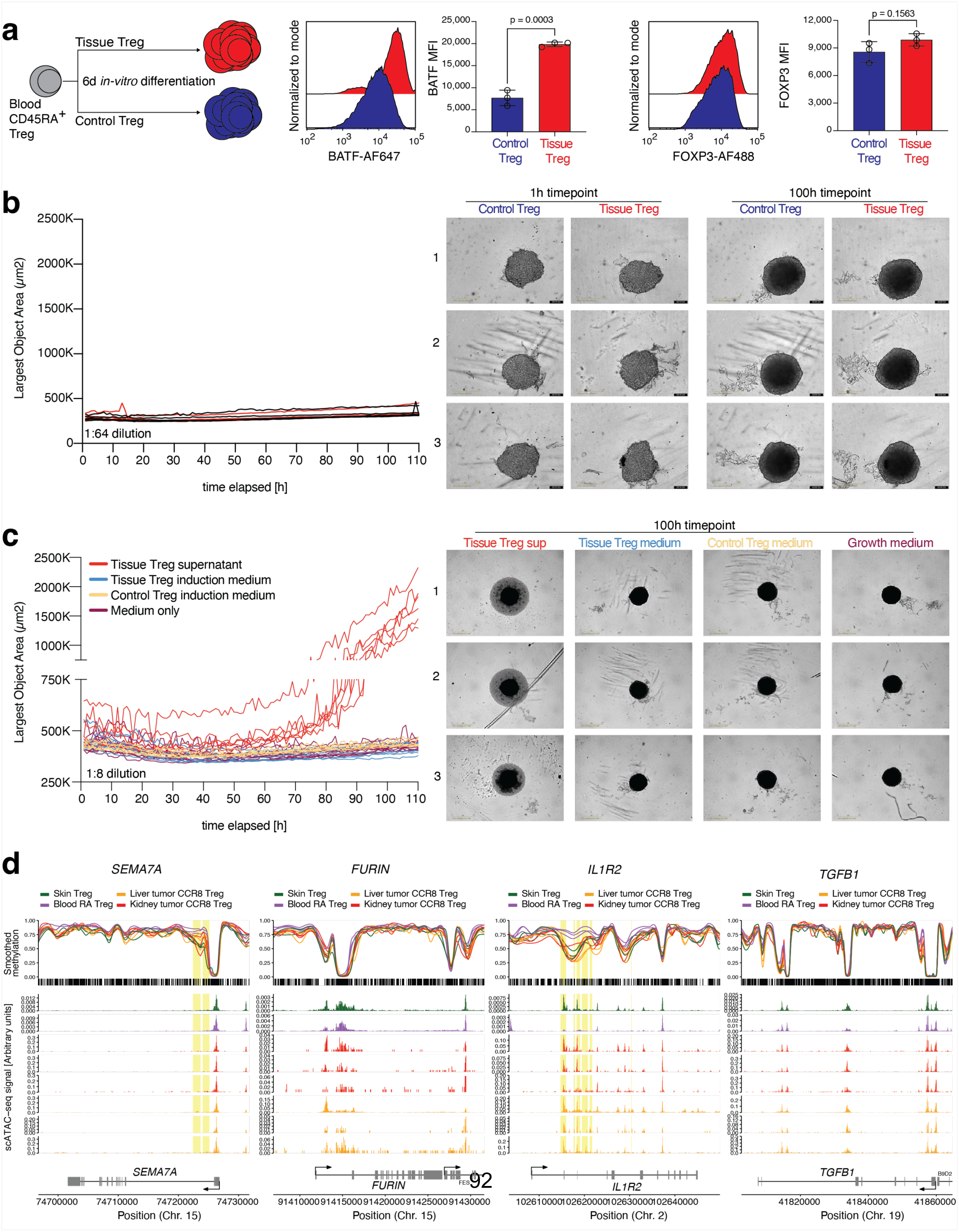
Controls for tumor spheroid cultures. **(a)** Induction of the tissue Treg-like program from human blood CD45RA^+^ Treg cells for 6 days *in vitro* using a stimulation and cytokine cocktail (TransAct, IL-2, TGF-β, IL-12, IL-21, IL-23) versus control Treg cells (TransAct, IL-2), results of intracellular FOXP3 and BATF staining (n=3, unpaired t test). **(b)** Tumor spheroids were seeded, treated with cell culture supernatant from either tissue Treg or control Treg cells (1:64 dilution) and imaged for 110 hours using the Incucyte Imaging chamber system. Representative images from 3 donors to the right (n=3 with two technical replicates). **(c)** Tumor spheroids were seeded, treated with cell culture supernatant from either tissue Treg medium (1:8 dilution, red lines), tissue Treg induction medium (IL-2, TGF-β, IL-12, IL-21, IL-23, blue lines), control Treg medium (IL-2, yellow lines) or serum-free growth medium only (red lines) and imaged for 110 hours using the Incucyte Imaging chamber system. Representative images from 3 donors to the right (n=3 with two technical replicates). **(d)** Smoothed methylation values for selected regions with blood CD45RA^+^ Treg, skin Treg, kidney tumor CCR8^+^ Treg and liver tumor CCR8^+^ Treg cells. Below, chromatin accessibility. Data representative of three or more independent experiments with three or more individual donors.

## REFERENCES

1. Sakaguchi, S. et al. Regulatory T Cells and Human Disease. Annu Rev Immunol 38, 541–566 (2020).

2. Burton, O.T. et al. The tissue-resident regulatory T cell pool is shaped by transient multi-tissue migration and a conserved residency program. Immunity 57, 1586–1602 e1510 (2024).

3. Ali, N. et al. Regulatory T Cells in Skin Facilitate Epithelial Stem Cell Differentiation. Cell 169, 1119–1129 e1111 (2017).

4. Shime, H. et al. Proenkephalin(+) regulatory T cells expanded by ultraviolet B exposure maintain skin homeostasis with a healing function. Proc Natl Acad Sci U S A 117, 20696–20705 (2020).

5. Nosbaum, A. et al. Cutting Edge: Regulatory T Cells Facilitate Cutaneous Wound Healing. J Immunol 196, 2010–2014 (2016).

6. Santamaria, E. et al. The Epidermal Growth Factor Receptor Ligand Amphiregulin Protects From Cholestatic Liver Injury and Regulates Bile Acids Synthesis. Hepatology 69, 1632–1647 (2019).

7. Feuerer, M. et al. Lean, but not obese, fat is enriched for a unique population of regulatory T cells that affect metabolic parameters. Nat Med 15, 930–939 (2009).

8. Sakai, R. et al. Kidney GATA3(+) regulatory T cells play roles in the convalescence stage after antibody-mediated renal injury. Cell Mol Immunol 18, 1249–1261 (2021).

9. Tan, W. et al. Interleukin-33-Dependent Accumulation of Regulatory T Cells Mediates Pulmonary Epithelial Regeneration During Acute Respiratory Distress Syndrome. Front Immunol 12, 653803 (2021).

10. Liu, Q., et al. IL-33-mediated IL-13 secretion by ST2+ Tregs controls inflammation after lung injury. JCI Insight 4 (2019).

11. Garibaldi, B.T. et al. Regulatory T cells reduce acute lung injury fibroproliferation by decreasing fibrocyte recruitment. Am J Respir Cell Mol Biol 48, 35–43 (2013).

12. Kaiser, K.A., Loffredo, L.F., Santos-Alexis, K.L., Ringham, O.R. & Arpaia, N. Regulation of the alveolar regenerative niche by amphiregulin-producing regulatory T cells. J Exp Med 220 (2023).

13. Arpaia, N. et al. A Distinct Function of Regulatory T Cells in Tissue Protection. Cell 162, 1078–1089 (2015).

14. Harb, H. et al. Notch4 signaling limits regulatory T-cell-mediated tissue repair and promotes severe lung inflammation in viral infections. Immunity 54, 1186–1199 e1187 (2021).

15. Burzyn, D. et al. A special population of regulatory T cells potentiates muscle repair. Cell 155, 1282–1295 (2013).

16. Kuswanto, W. et al. Poor Repair of Skeletal Muscle in Aging Mice Reflects a Defect in Local, Interleukin-33-Dependent Accumulation of Regulatory T Cells. Immunity 44, 355–367 (2016).

17. Xia, N. et al. A Unique Population of Regulatory T Cells in Heart Potentiates Cardiac Protection From Myocardial Infarction. Circulation 142, 1956–1973 (2020).

18. Zacchigna, S. et al. Paracrine effect of regulatory T cells promotes cardiomyocyte proliferation during pregnancy and after myocardial infarction. Nat Commun 9, 2432 (2018).

19. Weirather, J. et al. Foxp3+ CD4+ T cells improve healing after myocardial infarction by modulating monocyte/macrophage differentiation. Circ Res 115, 55–67 (2014).

20. Liesz, A. et al. Regulatory T cells are key cerebroprotective immunomodulators in acute experimental stroke. Nat Med 15, 192–199 (2009).

21. Ito, M. et al. Brain regulatory T cells suppress astrogliosis and potentiate neurological recovery. Nature 565, 246–250 (2019).

22. Dombrowski, Y. et al. Regulatory T cells promote myelin regeneration in the central nervous system. Nat Neurosci 20, 674–680 (2017).

23. Green, J.A., Arpaia, N., Schizas, M., Dobrin, A. & Rudensky, A.Y. A nonimmune function of T cells in promoting lung tumor progression. J Exp Med 214, 3565–3575 (2017).

24. Halvorsen, E.C. et al. IL-33 increases ST2(+) Tregs and promotes metastatic tumour growth in the lungs in an amphiregulin-dependent manner. Oncoimmunology 8, e1527497 (2019).

25. Wang, L. et al. AREG mediates the epithelial-mesenchymal transition in pancreatic cancer cells via the EGFR/ERK/NF-kappaB signalling pathway. Oncol Rep 43, 1558–1568 (2020).

26. Xu, Q. et al. Targeting amphiregulin (AREG) derived from senescent stromal cells diminishes cancer resistance and averts programmed cell death 1 ligand (PD-L1)-mediated immunosuppression. Aging Cell 18, e13027 (2019).

27. Braband, K.L. et al. Stepwise acquisition of unique epigenetic signatures during differentiation of tissue Treg cells. Front Immunol 13, 1082055 (2022).

28. Villarreal, D.O. et al. Targeting CCR8 Induces Protective Antitumor Immunity and Enhances Vaccine-Induced Responses in Colon Cancer. Cancer Res 78, 5340–5348 (2018).

29. Itahashi, K., et al. BATF epigenetically and transcriptionally controls the activation program of regulatory T cells in human tumors. Sci Immunol 7, eabk0957 (2022).

30. Van Damme, H. et al. Therapeutic depletion of CCR8(+) tumor-infiltrating regulatory T cells elicits antitumor immunity and synergizes with anti-PD-1 therapy. J Immunother Cancer 9 (2021).

31. Wang, T. et al. CCR8 blockade primes anti-tumor immunity through intratumoral regulatory T cells destabilization in muscle-invasive bladder cancer. Cancer Immunol Immunother 69, 1855–1867 (2020).

32. Plitas, G. et al. Regulatory T Cells Exhibit Distinct Features in Human Breast Cancer. Immunity 45, 1122–1134 (2016).

33. Kidani, Y. et al. CCR8-targeted specific depletion of clonally expanded Treg cells in tumor tissues evokes potent tumor immunity with long-lasting memory. Proc Natl Acad Sci U S A 119 (2022).

34. Liu, S. et al. CD4(+)CCR8(+) Tregs in ovarian cancer: a potential effector Tregs for immune regulation. J Transl Med 21, 803 (2023).

35. Delacher, M. et al. Single-cell chromatin accessibility landscape identifies tissue repair program in human regulatory T cells. Immunity 54, 702–720 e717 (2021).

36. Wang, Q. et al. Tagmentation-based whole-genome bisulfite sequencing. Nat Protoc 8, 2022–2032 (2013).

37. Weichenhan, D. et al. Tagmentation-Based Library Preparation for Low DNA Input Whole Genome Bisulfite Sequencing. Methods Mol Biol 1708, 105–122 (2018).

38. Beumer, N. et al. DNA hypomethylation comprising transposable elements defines human regulatory T cells in cutaneous tissue and identifies their blood recirculating counterpart. *Nat Immunol* **accepted in principle**, TBA (2025).

39. Ohkura, N. & Sakaguchi, S. Transcriptional and epigenetic basis of Treg cell development and function: its genetic anomalies or variations in autoimmune diseases. Cell Res 30, 465–474 (2020).

40. Kim, M. & Costello, J. DNA methylation: an epigenetic mark of cellular memory. Exp Mol Med 49, e322 (2017).

41. Pillarisetty, V.G. The pancreatic cancer microenvironment: an immunologic battleground. Oncoimmunology 3, e950171 (2014).

42. Braband, K.L. et al. Using single-cell chromatin accessibility sequencing to characterize CD4+ T cells from murine tissues. Front Immunol 14, 1232511 (2023).

43. Granja, J.M. et al. ArchR is a scalable software package for integrative single-cell chromatin accessibility analysis. Nat Genet 53, 403–411 (2021).

44. Delacher, M. et al. Genome-wide DNA-methylation landscape defines specialization of regulatory T cells in tissues. Nat Immunol 18, 1160–1172 (2017).

45. Delacher, M. et al. Precursors for Nonlymphoid-Tissue Treg Cells Reside in Secondary Lymphoid Organs and Are Programmed by the Transcription Factor BATF. Immunity 52, 295–312 e211 (2020).

46. Llovet, J.M. et al. Hepatocellular carcinoma. Nat Rev Dis Primers 2, 16018 (2016).

47. Liu, G., Xiong, D., Che, Z., Chen, H. & Jin, W. A novel inflammation-associated prognostic signature for clear cell renal cell carcinoma. Oncol Lett 24, 307 (2022).

48. Hovelmeyer, N., Schmidt-Supprian, M. & Ohnmacht, C. NF-kappaB in control of regulatory T cell development, identity, and function. J Mol Med (Berl) 100, 985–995 (2022).

49. Atsaves, V., Leventaki, V., Rassidakis, G.Z. & Claret, F.X. AP-1 Transcription Factors as Regulators of Immune Responses in Cancer. Cancers (Basel) 11 (2019).

50. Song, G. et al. TIMP1 is a prognostic marker for the progression and metastasis of colon cancer through FAK-PI3K/AKT and MAPK pathway. J Exp Clin Cancer Res 35, 148 (2016).

51. Pope, B.D. et al. Topologically associating domains are stable units of replication-timing regulation. Nature 515, 402–405 (2014).

52. Sadik, A. et al. IL4I1 Is a Metabolic Immune Checkpoint that Activates the AHR and Promotes Tumor Progression. Cell 182, 1252–1270 e1234 (2020).

53. Kennedy, P.T. et al. Soluble CTLA-4 attenuates T cell activation and modulates anti-tumor immunity. Mol Ther 32, 457–468 (2024).

54. Lecocq, Q., Keyaerts, M., Devoogdt, N. & Breckpot, K. The Next-Generation Immune Checkpoint LAG-3 and Its Therapeutic Potential in Oncology: Third Time’s a Charm. Int J Mol Sci 22 (2020).

55. Hargadon, K.M. Dysregulation of TGFbeta1 Activity in Cancer and Its Influence on the Quality of Anti-Tumor Immunity. J Clin Med 5 (2016).

56. Black, S.A., Nelson, A.C., Gurule, N.J., Futscher, B.W. & Lyons, T.R. Semaphorin 7a exerts pleiotropic effects to promote breast tumor progression. Oncogene 35, 5170–5178 (2016).

57. Pesu, M. et al. T-cell-expressed proprotein convertase furin is essential for maintenance of peripheral immune tolerance. Nature 455, 246–250 (2008).

58. Guo, H. et al. Single-cell RNA sequencing reveals an IL1R2+Treg subset driving immunosuppressive microenvironment in HNSCC. Cancer Immunol Immunother 74, 159 (2025).

59. De Simone, M. et al. Transcriptional Landscape of Human Tissue Lymphocytes Unveils Uniqueness of Tumor-Infiltrating T Regulatory Cells. Immunity 45, 1135–1147 (2016).

60. Alvisi, G. et al. IRF4 instructs effector Treg differentiation and immune suppression in human cancer. J Clin Invest 130, 3137–3150 (2020).

61. Weaver, J.D. et al. Differential expression of CCR8 in tumors versus normal tissue allows specific depletion of tumor-infiltrating T regulatory cells by GS-1811, a novel Fc-optimized anti-CCR8 antibody. Oncoimmunology 11, 2141007 (2022).

62. Mijnheer, G. et al. Conserved human effector Treg cell transcriptomic and epigenetic signature in arthritic joint inflammation. Nat Commun 12, 2710 (2021).

63. Zhang, Y. et al. Model-based analysis of ChIP-Seq (MACS). Genome Biol 9, R137 (2008).

64. Chung, N.C., Miasojedow, B., Startek, M. & Gambin, A. Jaccard/Tanimoto similarity test and estimation methods for biological presence-absence data. BMC Bioinformatics 20, 644 (2019).

65. Zhao, H. et al. CrossMap: a versatile tool for coordinate conversion between genome assemblies. Bioinformatics 30, 1006–1007 (2014).

66. Stuart, T., Srivastava, A., Madad, S., Lareau, C.A. & Satija, R. Single-cell chromatin state analysis with Signac. Nat Methods 18, 1333–1341 (2021).

67. Satija, R., Farrell, J.A., Gennert, D., Schier, A.F. & Regev, A. Spatial reconstruction of single-cell gene expression data. Nat Biotechnol 33, 495–502 (2015).

68. Stuart, T. et al. Comprehensive Integration of Single-Cell Data. Cell 177, 1888–1902 e1821 (2019).

69. Gu, Z., Eils, R. & Schlesner, M. Complex heatmaps reveal patterns and correlations in multidimensional genomic data. Bioinformatics 32, 2847–2849 (2016).

70. Hovestadt, V. et al. Decoding the regulatory landscape of medulloblastoma using DNA methylation sequencing. Nature 510, 537–541 (2014).

71. Bolger, A.M., Lohse, M. & Usadel, B. Trimmomatic: a flexible trimmer for Illumina sequence data. Bioinformatics 30, 2114–2120 (2014).

72. Li, H. & Durbin, R. Fast and accurate short read alignment with Burrows-Wheeler transform. Bioinformatics 25, 1754–1760 (2009).

73. Tarasov, A., Vilella, A.J., Cuppen, E., Nijman, I.J. & Prins, P. Sambamba: fast processing of NGS alignment formats. Bioinformatics 31, 2032–2034 (2015).

74. Hansen, K.D., Langmead, B. & Irizarry, R.A. BSmooth: from whole genome bisulfite sequencing reads to differentially methylated regions. Genome Biol 13, R83 (2012).

75. Gu, Z., Gu, L., Eils, R., Schlesner, M. & Brors, B. circlize Implements and enhances circular visualization in R. Bioinformatics 30, 2811–2812 (2014).

76. Ramirez, F. et al. deepTools2: a next generation web server for deep-sequencing data analysis. Nucleic Acids Res 44, W160–165 (2016).

77. Lawrence, M., Gentleman, R. & Carey, V. rtracklayer: an R package for interfacing with genome browsers. Bioinformatics 25, 1841–1842 (2009).

78. Frankish, A. et al. GENCODE reference annotation for the human and mouse genomes. Nucleic Acids Res 47, D766–D773 (2019).

79. Hughes, C.S. et al. Ultrasensitive proteome analysis using paramagnetic bead technology. Mol Syst Biol 10, 757 (2014).

80. Sielaff, M. et al. Evaluation of FASP, SP3, and iST Protocols for Proteomic Sample Preparation in the Low Microgram Range. J Proteome Res 16, 4060–4072 (2017).

81. Wisniewski, J.R., Zougman, A., Nagaraj, N. & Mann, M. Universal sample preparation method for proteome analysis. Nat Methods 6, 359–362 (2009).

82. Distler, U., Kuharev, J., Navarro, P. & Tenzer, S. Label-free quantification in ion mobility-enhanced data-independent acquisition proteomics. Nat Protoc 11, 795–812 (2016).

83. Meier, F. et al. diaPASEF: parallel accumulation-serial fragmentation combined with data-independent acquisition. Nat Methods 17, 1229–1236 (2020).

84. Skowronek, P. et al. Rapid and In-Depth Coverage of the (Phospho-)Proteome With Deep Libraries and Optimal Window Design for dia-PASEF. Mol Cell Proteomics 21, 100279 (2022).

85. Demichev, V., Messner, C.B., Vernardis, S.I., Lilley, K.S. & Ralser, M. DIA-NN: neural networks and interference correction enable deep proteome coverage in high throughput. Nat Methods 17, 41–44 (2020).

86. Muller-Dott, S. et al. Expanding the coverage of regulons from high-confidence prior knowledge for accurate estimation of transcription factor activities. Nucleic Acids Res 51, 10934–10949 (2023).

87. Badia, I.M.P., et al. decoupleR: ensemble of computational methods to infer biological activities from omics data. Bioinform Adv 2, vbac016 (2022).

